# Molecular Mimicry Map (3M) of SARS-CoV-2: Prediction of potentially immunopathogenic SARS-CoV-2 epitopes via a novel immunoinformatic approach

**DOI:** 10.1101/2020.11.12.344424

**Authors:** Hyunsu An, Jihwan Park

**Author notes:** **Correspondence**, Jihwan Park, PhD., Assistant Professor, School of Life Sciences, Gwangju Institute of Science and Technology (GIST), 123 Cheomdangwagi-ro, Buk-gu, Gwangju, Republic of Korea.

## Abstract

Currently, more than 33 million peoples have been infected by severe acute respiratory syndrome coronavirus 2 (SARS-CoV-2), and more than a million people died from coronavirus disease 2019 (COVID-19), a disease caused by the virus. There have been multiple reports of autoimmune and inflammatory diseases following SARS-CoV-2 infections. There are several suggested mechanisms involved in the development of autoimmune diseases, including cross-reactivity (molecular mimicry). A typical workflow for discovering cross-reactive epitopes (mimotopes) starts with a sequence similarity search between protein sequences of human and a pathogen. However, sequence similarity information alone is not enough to predict cross-reactivity between proteins since proteins can share highly similar conformational epitopes whose amino acid residues are situated far apart in the linear protein sequences. Therefore, we used a hidden Markov model-based tool to identify distant viral homologs of human proteins. Also, we utilized experimentally determined and modeled protein structures of SARS-CoV-2 and human proteins to find homologous protein structures between them. Next, we predicted binding affinity (IC50) of potentially cross-reactive T-cell epitopes to 34 MHC allelic variants that have been associated with autoimmune diseases using multiple prediction algorithms. Overall, from 8,138 SARS-CoV-2 genomes, we identified 3,238 potentially cross-reactive B-cell epitopes covering six human proteins and 1,224 potentially cross-reactive T-cell epitopes covering 285 human proteins. To visualize the predicted cross-reactive T-cell and B-cell epitopes, we developed a web-based application “Molecular Mimicry Map (3M) of SARS-CoV-2” (available at https://ahs2202.github.io/3M/). The web application enables researchers to explore potential cross-reactive SARS-CoV-2 epitopes alongside custom peptide vaccines, allowing researchers to identify potentially suboptimal peptide vaccine candidates or less ideal part of a whole virus vaccine to design a safer vaccine for people with genetic and environmental predispositions to autoimmune diseases. Together, the computational resources and the interactive web application provide a foundation for the investigation of molecular mimicry in the pathogenesis of autoimmune disease following COVID-19.

## Introduction

Since February 2020, severe acute respiratory syndrome coronavirus 2 (SARS-CoV-2) has been spreading across every continent except Antarctica, affecting virtually all nations in the world. Currently, more than 45 million peoples have been infected by SARS-CoV-2, and more than a million people died from coronavirus disease 2019 (COVID-19), a disease caused by the virus.

Critical medical conditions associated with severe cases of COVID-19 include acute respiratory distress syndrome (ARDS), cytokine release syndrome (CRS), multiorgan dysfunction syndrome, and disseminated intravascular coagulation (DIC). Endothelial dysfunction and extensive damage to endothelial cells due to systemic inflammatory response have been attributed to the pathology of DIC associated with COVID-19 (Merrill et al., 2020).

There have been multiple reports of autoimmune and inflammatory diseases following SARS-CoV-2 infections. These autoimmune diseases include Kawasaki-like multisystem inflammatory syndrome in children (MIS-C) (Akca et al., 2020; Gruber et al., 2020; Verdoni et al., 2020; Whittaker et al., 2020), Guillain–Barré syndrome (GBS) (Ottaviani et al., 2020; Schiaffino et al., 2020; Sedaghat and Karimi, 2020), myositis (Mehan et al., 2020; Zhang et al., 2020a), IgA vasculitis with nephritis (Su et al., 2020; Suso et al., 2020), membranous glomerulonephritis (MGN) (Kudose et al., 2020), antineutrophil cytoplasmic antibodies (ANCA) associated vasculitis (Uppal et al., 2020), autoimmune hemolytic anemia (AIHA) (Jensen et al., 2020; Lopez et al., 2020), antiphospholipid syndrome (APS) (Amezcua-Guerra et al., 2020; Zhang et al., 2020b), autoimmune thrombotic thrombocytopenic purpura (TTP) (Albiol et al., 2020; Hindilerden et al., 2020), and immune thrombocytopenic purpura (ITP) (Kewan et al., 2020; Tsao et al., 2020; Zulfiqar et al., 2020). Even though there is no direct evidence that SARS-CoV-2 infection caused the series of immunopathological reactions that eventually caused the autoimmune diseases, it appears that there is enough circumstantial evidence to suspect that the autoimmune responses were caused by, or at least initiated by, our body’s response to SARS-CoV-2 infection.

In the current model of autoimmune diseases, both genetics and environment such as microbiota play roles in the loss of self-tolerance, development of autoreactive B-cells and T-cells, and the pathogenesis of autoimmune diseases (Rojas et al., 2018; Ruff et al., 2020). There are several suggested mechanisms involved in the development of autoimmune diseases, including bystander activation (Kim and Shin, 2019; McCoy et al., 2006), cross-reactivity of antibodies (B-cell cross-reactivity) (Luo et al., 2018b; McCoy et al., 2006; Wucherpfennig and Strominger, 1995; Zhao et al., 1998), T-cell receptor degeneracy (T-cell receptor cross-reactivity) (Cole et al., 2016; Riley et al., 2018), dual T-cell receptors (Ji et al., 2010), and epitope spreading (Miller et al., 1997; Seitz-Polski et al., 2018; Sokolove et al., 2012). Molecular mimicry refers to the ability of epitopes derived from bacterial and viral pathogens to activate autoreactive T-cells and B-cells. A term mimotope describes an epitope from invading pathogens that mimic an exposed surface of a folded human protein (self B-cell epitopes), the major histocompatibility complex (MHC) molecule-bound peptides derived from endogenous human proteins (self T-cell epitopes), carbohydrate, or lipid antigen present in human cells. T-cell receptor cross-reactivity describes a specific subset of T-cell receptors capable of recognizing multiple MHC-presented peptides, which can be as diverse as 1 million different peptides in the case of 1E6 CD8+ T cell clone (Cole et al., 2016). Also, very divergent epitopes without significant sequence similarity can bind the same T-cell receptor (TCR) by assuming different conformations when binding the TCR (Riley et al., 2018). Unlike a folded protein with a rigid structure (common for B-cell epitopes), peptides (T-cell epitopes) are more flexible and can assume diverse conformations when binding T-cell receptors (TCRs). During infection, potentially cross-reactive epitopes from an invading pathogen can initiate a series of pathological immune reactions leading to the development of autoimmune diseases. One of the most classical examples of infection-induced autoimmunity is rheumatic fever (RF) caused by group A streptococcal infection of the upper respiratory tract (Tandon et al., 2013). In rheumatic fever, the sequence similarity of streptococcal M proteins to the basement membrane collagen (type IV) leads to the development of autoantibodies directed against collagen type IV proteins, which can cause acute inflammation of the heart (carditis) and permanent damage to heart valves (rheumatic heart disease, RHD) (Tandon et al., 2013). There has been an increasing number of investigations of the involvement of molecular mimicry in the roles of microbiome in the development of autoimmune diseases (Gkoutzourelas et al., 2020; Gourh et al., 2020; Greiling et al., 2018; Petersen et al., 2020) or the pathogenesis of autoimmune symptoms following bacterial or viral infections (Amedei et al., 2003; Chuang et al., 2014; Luo et al., 2018b; Venigalla et al., 2020) or vaccination (Luo et al., 2018b).

Activation of cross-reactive T-cells and B-cells (molecular mimicry) combined with impaired immune regulation in COVID-19 patients can play a role in the immunopathogenesis of severe COVID-19 symptoms and autoimmune diseases following SARS-CoV-2 infection. It has been suggested that the combined action of molecular mimicry and bystander activation can contribute to the pathogenesis of autoimmune diseases (McCoy et al., 2006). Autoreactive T-cells and B-cells that escaped central tolerance in the thymus and the bone marrow, respectively, are further suppressed via peripheral immunological tolerance, where tolerogenic dendritic cells (tolDCs) and regulatory T cells (Tregs) suppress the activation of autoreactive T-cells and B-cells. A decreased number of regulatory T-cells has been associated with severe COVID-19 cases (Qin et al., 2020), which implies that mechanisms that mediate peripheral tolerance are dysfunctional in severe COVID-19 patients. Moreover, the inflammatory milieu created in the lungs and increased cytokines in the blood of severe COVID-19 patients can further activate autoreactive T-cells and B-cells that have escaped the central tolerance mechanism via bystander activation. Recent deep immune profiling of hospitalized COVID-19 patients revealed broad activation of all non-naïve cytotoxic CD8+ T cells in ~80% of hospitalized COVID-19 patients, which was correlated with elevations of inflammation markers including IL-6 and ferritin levels (Mathew et al., 2020).

A couple of recent reviews explored the possibility of involvement of molecular mimicry in the immunopathology observed in severe COVID-19 patients (Cappello et al., 2020; Rodriguez et al., 2020; Shah et al., 2020; Talotta and Robertson, 2020). Molecular mimicry has been proposed as an underlying cause of serious complications of COVID-19 such as disseminated intravascular coagulation (DIC) due to damage to endothelial cells (Marino Gammazza et al., 2020), respiratory failure due to damage to the respiratory pacemaker in the brainstem (pre-Bötzinger complex) (Lucchese and Floel, 2020a), and multiorgan dysfunction syndrome (Angileri et al., 2020a; Cappello, 2020a, b), autoimmune hemolytic anemia (Angileri et al., 2020b), Guillain-Barré syndrome (Lucchese and Floel, 2020b; Schiaffino et al., 2020), and other complications (Kanduc, 2020; Lucchese and Floel, 2020b; Megremis et al., 2020). For example, Schiaffino and colleagues reported the presence of autoantibodies targeting liver and gut mucosa in about of a quarter of hospitalized COVID-19 patients (12 of 52 patients) in a small cohort (Schiaffino et al., 2020). They reported a novel immunofluorescence pattern (COVID-IF pattern) of cross-reactive autoantibodies in the serums of twelve COVID-19 patients in the cohort of 52 COVID-19 patients (Schiaffino et al., 2020). Using the rat tissues, the authors detected the presence of cross-reactive autoantibodies in the serum of COVID-19 patients, which reacted with antigens on the plasma membrane of hepatocytes and gastric mucosa cells. Interestingly, three patients with high titer of COVID-19 specific autoantibody in their serum also had severe neurological complications from COVID-19. The cerebrospinal fluid (CSF) of one of the three patients also produced the COVID-19 associated immunofluorescence pattern, who also had Guillain–Barré syndrome. Antinuclear antibodies (ANA) were not detected in serums of the twelve COVID-19 patients with the novel immunofluorescence staining pattern.

Based on current knowledge of how molecular mimicry can initiate the pathology of autoimmune disease, we designed a novel immunoinformatic approach to identify potentially cross-reactive B-cell and T-cell epitopes with increased sensitivity and accuracy. We first assigned structural properties of each residue in the human proteome and thousands of non-redundant SARS-CoV-2 using experimentally determined and modeled protein structures. We then used our structural property database of human and SARS-CoV-2 proteomes to find shared conformational epitopes (B-cell epitopes) of human and SARS-CoV-2 proteins. We also predicted binding affinity (IC50) of a pair of aligned peptides from human and SARS-CoV-2 proteins to autoimmune disease-associated MHC allelic variants to identify potentially cross-reactive MHC-I and MHC-II ligands. To visualize the predicted cross-reactive T-cell and B-cell epitopes of SARS-CoV-2 proteins, we developed a web-based application “Molecular Mimicry Map of SARS-CoV-2” or “3M of SARS-CoV-2” (available at https://ahs2202.github.io/3M/).

## Results

### Alignment between human proteome and 8,138 SARS-CoV-2 proteomes

A typical workflow for the discovery of mimotopes starts with a sequence similarity search between human protein sequences and protein sequences of the pathogen. Due to its ease of use and wide availability, Basic Local Alignment Search Tool (BLAST) from NCBI (Camacho et al., 2009) is commonly used to find a stretch of amino acids in bacterial or viral proteins that harbor significant similarity to human proteins (Gkoutzourelas et al., 2020; Greiling et al., 2018; Polymeros et al., 2014; Venigalla et al., 2020). Protein BLAST (BLASTP) has also been utilized to find potential mimotopes of SARS-CoV-2 virus proteins to suggests a possible involvement of molecular mimicry in the pathogenesis of the neurological symptoms from COVID-19 (Lucchese and Floel, 2020a) and autoimmune diseases following SARS-CoV-2 infections (Angileri et al., 2020b; Megremis et al., 2020).

BLASTP can efficiently and accurately identify similar regions between two nucleotide or protein sequences. BLASTP utilizes the BLOSUM62 scoring matrix to calculate a similarity score between two aligned protein sequences. Since a single scoring matrix is used to calculate the similarity score for every position in the alignment, a distantly related sequence with frequent insertions, deletions, and non-conservative substitutions is less likely to be detected when the BLASTP algorithm is used alone. Therefore, it is less equipped to detect homologous proteins with low overall sequence similarity, such as bacterial or viral proteins that are remotely homologous to human proteins. However, even with low overall sequence similarity, proteins can share highly similar surface epitopes, which often consists of a collection of discontinuous patches of amino acids that are often situated far apart in the protein sequences. Therefore, it is important to identify even very distant bacterial or viral homologs of human proteins to sensitively detect shared surface epitopes between viral proteins and human proteins. To detect remote viral homologs of human proteins in SARS-CoV-2 proteomes with increased sensitivity, HMMER (Eddy, 2011) was utilized to build hidden Markov models of viral homologs of human proteins. HMMER learns a position-specific scoring matrix tailored for each position of a given multiple sequence alignment (MSA) of viral homologs of a human protein to detect a remote homology between SARS-CoV-2 and human proteins. To identify viral homologs of human proteins, we iteratively searched 2,949,581 viral proteins of all human viruses whose protein sequences are available in the NCBI Virus resource (accessed 26 June 2020) with every human protein sequence. We created a database of profile hidden Markov models (profile HMM) from a collection of multiple sequence alignments between each human protein sequence and viral protein sequences containing a homologous region to the human protein. We then used the database to identify regions of SARS-CoV-2 proteins that share significant homology with human protein sequences.

Overall, 93,413 viral protein sequences from 8,138 SARS-CoV-2 proteomes were searched against 20,595 human protein sequences using BLASTP and HMMER tools. The sequence similarity search results from BLASTP and HMMER were combined, resulting in 1,622,641 alignments between SARS-CoV-2 and human protein sequences. After the deduplication of redundant SARS-CoV-2 protein sequences, 150,744 alignments between non-redundant SARS-CoV-2 protein sequences and human proteins were further analyzed to predict potentially cross-reactive B-cell epitopes and T-cell epitopes.

### Prediction of B-Cell epitopes of SARS-CoV-2 that can potentially induce type II and III hypersensitivities using a novel immunoinformatic approach

Next, we utilized experimentally determined and modeled protein structures of SARS-CoV-2 virus and human proteins to find homologous protein structures between SARS-CoV-2 and human. We then identified amino acid residues accessible on the surface of the homologous protein structures and thus accessible to circulating antibodies. We reasoned that such co-accessible regions of homologous protein structures of SARS-CoV-2 and human can share highly similar conformational epitopes that can be recognized by the same antibody, representing potentially cross-reactive B-cell epitopes. Type II hypersensitivity classification of autoimmune reactions refers to an antibody-mediated process where autoreactive IgG and IgM antibodies bind autoantigens on cells or extracellular matrix, leading to tissue or organ-specific damage. Type III hypersensitivity is caused by the deposition of circulating immune complex to vascular walls and the glomerulus of the kidney, often affecting multiple organs. Antibodies implicated in type III hypersensitivity binds small soluble proteins including circulating bacterial or viral antigens during active infection (reactive arthritis) and autoantigens (systemic lupus erythematosus), that are normally sequestered inside a cell but exposed to extracellular space by the release of a neutrophil extracellular trap (NETosis), rupture of damaged cells (necrosis), or inappropriate clearance of apoptotic debris. Considering the possibility that proteins that are not normally exposed to the cell surface can be released to extracellular space during intense tissue damage, we also predicted potentially cross-reactive B-cell epitopes of non-structural proteins of SARS-CoV-2 as well as structural proteins of the virus.

To identify which amino acid residues are exposed to the surface of a protein and are capable of interacting with an antibody, we used experimentally determined protein structures from the Protein Data Bank (PDB) to assign relative surface availability (RSA) to each residue in the protein. For SARS-CoV-2 virus proteins, we additionally used template-based protein structure modeling results (Heo and Feig, 2020; Huang et al., 2020; Waterhouse et al., 2018) to assign RSA and other structural properties to residues whose structural information is not available in the PDB database (**Fig. S1**). After assigning structural properties to a protein using the PDB database and protein structure modeling results, the SCRATCH protein structural property predictor (Magnan and Baldi, 2014) was used to predict relative surface availability and secondary structure classification of remaining residues without assigned structural information (**Fig. S2**). The number of residues that were covered by each method for each protein of the SARS-CoV-2 reference proteome is given in **Table 1**. As a result, for a window size of 30 amino acids, we found 6,079 potentially cross-reactive B-cell epitopes (E-value < 0.05) between 23 human proteins and 165 non-redundant SARS-CoV-2 virus proteins from 8,138 SARS-CoV-2 genomes.

**Table 1.** The number of residues for each method that was used to assign a relative surface accessibility value to a residue.

|  | Number of residues with relative surface accessibility |  |  |
| --- | --- | --- | --- |
|  | Predicted | Modeling | Experimentally determined |
| ORF1ab polyprotein | 2 | 3160 | 3935 |
| surface glycoprotein (S) | 0 | 84 | 1189 |
| ORF3a protein | 0 | 83 | 193 |
| envelope protein (E) | 0 | 17 | 58 |
| membrane glycoprotein (M) | 0 | 222 | 0 |
| ORF6 protein | 3 | 60 | 0 |
| ORF7a protein | 0 | 37 | 84 |
| ORF7b protein | 0 | 43 | 0 |
| ORF8 protein | 5 | 120 | 0 |
| nucleocapsid phosphoprotein (N) | 0 | 157 | 262 |
| ORF10 protein | 0 | 38 | 0 |

Interestingly, we discovered significant sequence and structural similarities between the Macrodomain of SARS-CoV-2 nonstructural protein 3 (nsp3, ORF1ab from 819 to 2763) and several human enzyme proteins involved in ADP-ribosylation of proteins. These human enzymes include ADP-ribose glycohydrolase MACROD1, MACROD2, Protein mono-ADP-ribosyltransferase PARP14, and PARP9 (**Fig. 1**). We also detect less significant similarities between potential conformational epitopes of SARS-CoV-2 nonstructural protein 13 (nsp13, ORF1ab from 5325 to 5925) and the following five human proteins: helicase MOV-10 (MOV10), RNA helicase aquarius (AQR), probable helicase senataxin (SETX), DNA-binding protein SMUBP-2 (IGHMBP2), and Protein ZGRF1. Although we observed significant structural similarity between the Helicase domain of SARS-CoV-2 nsp13 and the human proteins, the pairs show relatively lower similarities of accessible amino acid residues exposed on the surface of the proteins. Other than the Macrodomain of nsp3 and the Helicase domain of nsp13, no other parts of the SARS-CoV-2 proteome harbored significant similarity (alignment identity > 50% for a window size of 100 amino acids).

**Figure 1.**
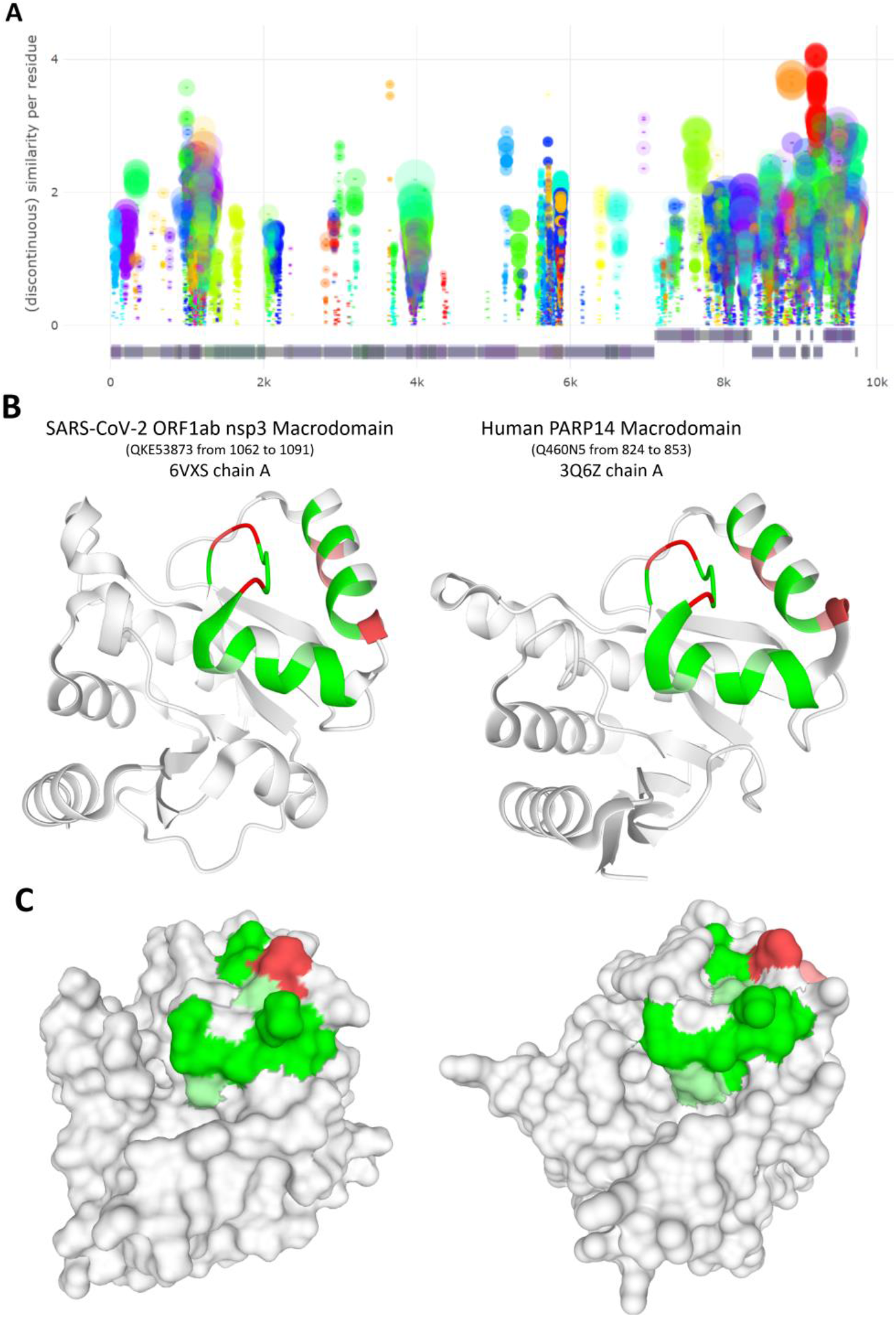
Identification of potentially cross-reactive B-cell epitopes of SARS-CoV-2 and human proteins. (A) Potentially cross-reactive B-cell epitopes of SARS-CoV-2 and human proteins visualized on SARS-CoV-2 representative proteome (window length is 30 amino acids). The size of the bubble represents the predicted cross-reactivity score between the SARS-CoV-2 epitope and the human protein, while the opacity of the bubble represents the correlation coefficient of relative surface availability values of the pair of cross-reactive epitopes of SARS-CoV-2 and human proteins. The color of the epitopes represents the name of a human protein from which the epitope was derived. (B, C) Experimentally determined structures of (left) Macrodomain of SARS-CoV-2 nonstructural protein 3 (nsp3) (PDB ID: 6VXS, Chain A) and (right) human PARP14 (PDB ID: 3Q6Z, Chain A).

### Prediction of T-cell epitopes of SARS-CoV-2 that can potentially induce type IV hypersensitivity using a novel immunoinformatic approach

Unlike type II and type III hypersensitivity, type IV hypersensitivity is mediated by CD4+ and CD8+ T-cells without the involvement of an antibody response. Type IV hypersensitivity is mediated by CD4+ type 1 helper T cells (Th1) and CD8+ cytotoxic T cells, and it is classified as a delayed-type hypersensitivity since the response requires more than 12 hours after exposure to antigens. There have been increasing efforts for identifying cross-reactive T-cell epitopes of pathogens that can initiate type IV hypersensitivity reactions and contribute to the pathogenesis of autoimmune diseases via activation of autoreactive CD8+ cytotoxic T cells (Luo et al., 2018b; Venigalla et al., 2020; Whalley et al., 2020; Yin et al., 2020).

As suggested by Luo et al. (Luo et al., 2018b), molecular mimicry can initiate an autoimmune reaction without the involvement of autoantibody. Luo and colleagues found that a subset of CD4+ T cells from type 1 narcolepsy (T1N) patients positive for MHC-II DQB1*06:02 allele can be stimulated by either peptide derived from 2009 H1N1 virus or peptides derived from HCRT and RFX4 proteins abundant in the hypocretin/orexin (HCRT) neuron. They suggested that the presence of autoreactive CD4+ T cells in T1N might be responsible for the destruction of the HCRT neurons by CD8+ T cells without the involvement of autoreactive antibodies, which is rarely detected in T1N. T1N has been associated with MHC-II DQB1*06:02 allele. Like T1N, autoimmune diseases with possible involvement of type IV hypersensitivity reactions have been associated with specific MHC-I or MHC-II alleles (Matzaraki et al., 2017). Therefore, we searched potentially cross-reactive MHC-I or MHC-II ligands of SARS-CoV-2 and human proteins for 34 autoimmune-associated MHC alleles (18 MHC-I alleles and 16 MHC-II alleles) (Matzaraki et al., 2017) using pVACtools (Hundal et al., 2020), an immunoinformatic toolkit for predicting MHC ligands using a variety of prediction algorithms. MHC class I molecules are loaded with peptides with 8-12 amino acids largely derived from intracellular proteins digested by the proteasome, while MHC class II molecules are loaded with peptides with 12-18 amino acids derived from extracellular proteins proteolytically digested inside a late endosome of antigen-presenting cells (Khodadoust et al., 2017; Wieczorek et al., 2017).

To predict potentially cross-reactive T-cell epitopes of SARS-CoV-2 virus and human proteins, we used predicted binding affinity (IC50) of potentially cross-reactive T-cell epitopes to MHC molecules associated with autoimmune diseases (Matzaraki et al., 2017). To increase the robustness of our prediction, we calculated geometric averages of predicted IC50 (nM) values from multiple prediction algorithms including MHCflurry (O’Donnell et al., 2020), MHCnuggets (Shao et al., 2020), NetMHC (Andreatta and Nielsen, 2016), NetMHCcons (Karosiene et al., 2012), NetMHCpan (Jurtz et al., 2017), NetMHCIIpan (Reynisson et al., 2020), NNalign (Nielsen and Andreatta, 2017), SMMalign (Nielsen et al., 2007), PickPocket (Karosiene et al., 2012), and SMMPMBEC (Kim et al., 2009). We retrieved 23,921, 27,230, 23,577, and 17,557 pairs of aligned human and virus peptides of 8, 9, 10, 15 amino acids in lengths, and predicted their binding affinities for 34 autoimmune-associated MHC alleles using the consensus of a maximum of eight algorithms (**Fig. S3**). Overall, from 8,138 SARS-CoV-2 proteomes, we identified 1,224 potentially cross-reactive T-cell epitopes (geometric average of predicted IC50 values < 100nM for both human and virus peptides) covering 285 human proteins after reducing redundancy and stringent filtering.

As of August 2020, there are six recombinant SARS-CoV-2 Spike protein-based vaccines undergoing Phase I or Phase II trials, and more than forty Spike protein-based vaccines in their preclinical phase (Krammer, 2020). A couple of recent investigations reported shared hexapeptide and heptapeptide sequences between SARS-CoV-2 Spike protein and human proteins (Kanduc, 2020; Kanduc and Shoenfeld, 2020). However, these studies did not evaluate whether the shared short peptide sequences between human and virus proteins can also be cross-reactive MHC ligands. After stringent filtering with predicted IC50 values less than 250nM and an average BLOSUM62 score per residue greater than 2.75, we identified 20 potentially cross-reactive MHC ligands of SARS-CoV-2 Spike protein and several human proteins (**Fig. 2**). Six potentially cross-reactive T-cell epitopes with average BLOSUM62 score per residue greater than 3.0 are listed in **Table 2**.

**Figure 2.**
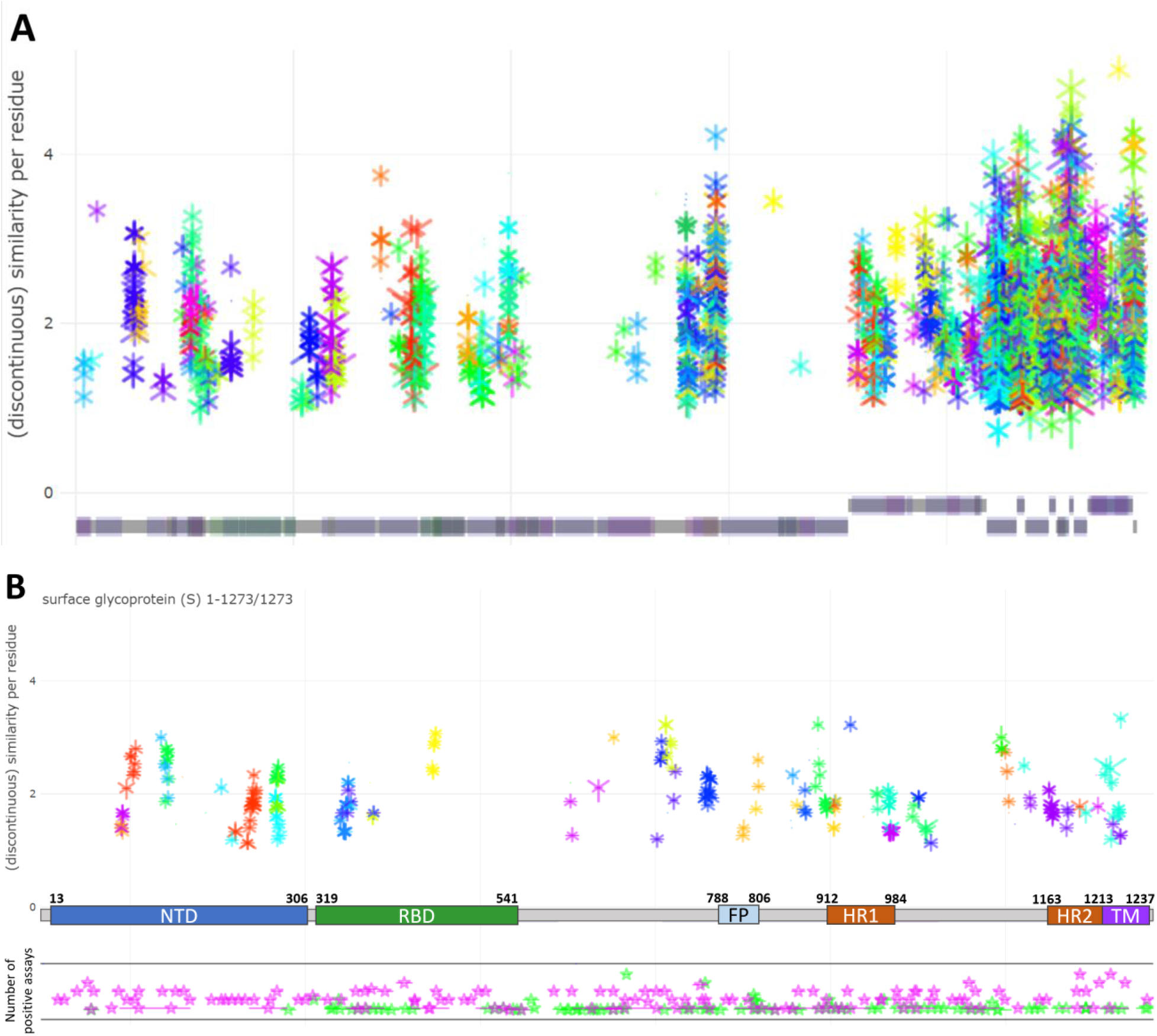
Identification of potentially cross-reactive T-cell epitopes of SARS-CoV-2 and human proteins. (A) Potentially cross-reactive T-cell epitopes of SARS-CoV-2 and human proteins visualized on SARS-CoV-2 representative proteome. A potentially cross-reactive T-cell epitope is represented by a line with a Y-shaped symbol and an upside-down-Y-shaped symbol in the middle of the line, representing crossreactive epitopes of human protein and SARS-CoV-2 protein, respectively. The start and end of the line represent the aligned position of the epitope on the representative SARS-CoV-2 protein. The size of either Y-shaped or upside-down-Y-shaped symbol represents a negative log (base 10) of the predicted binding affinity (IC50 in nM) of the epitope to the particular allelic variant of an MHC molecule. The color of the epitopes represents the name of a human protein from which the epitope was derived. (B) Potentially crossreactive T-cell epitopes of SARS-CoV-2 spike glycoprotein (S) and human proteins after stringent filtering with predicted binding affinity (IC50) to the given MHC allelic variant < 250nM for both SARS-CoV-2 and human epitopes. Experimentally validated SARS-CoV-1 and SARS-CoV-2 epitopes aligned to representative SARS-CoV-2 Spike protein is shown below the epitopes. Pink and green stars represent epitopes with positive T-cell and B-cell assays, respectively.

**Table 2.**
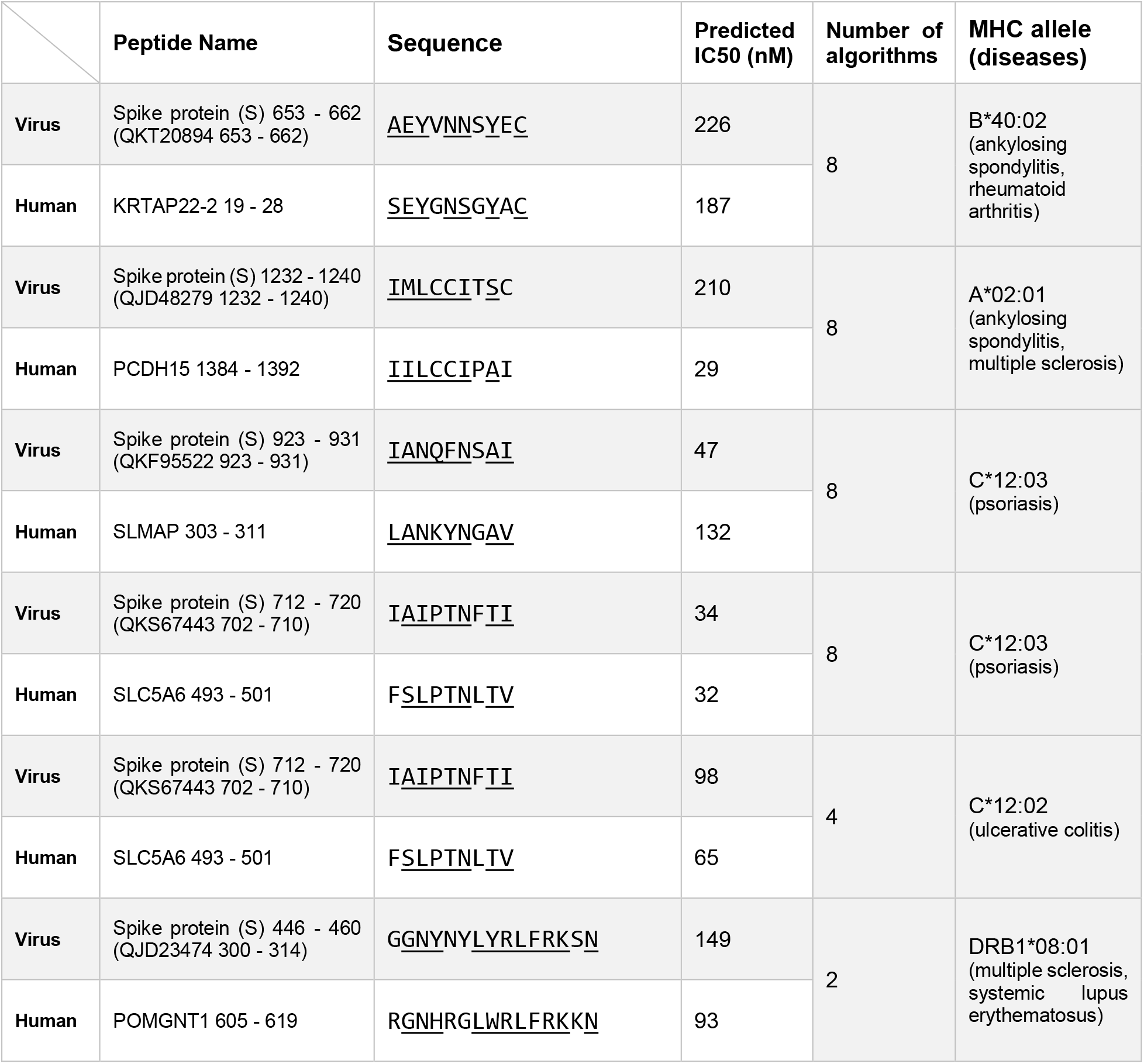
Identified potentially cross-reactive T-cell epitopes of SARS-CoV-2 Spike protein and human proteins

Molecular mimicry between human and bacterial heat shock proteins and the development of autoantibodies against heat shock proteins have been suggested to be involved in the initiation and pathogenesis of various inflammatory diseases, including atherosclerosis (Mandal et al., 2004; Zhu et al., 2001). Recently, molecular mimicry between human heat shock proteins and SARS-CoV-2 proteins has been suggested to be a potential pathogenic mechanism for extensive damage to endothelial cells in COVID-19 patients (Marino Gammazza et al., 2020) and Guillain-Barré syndrome following COVID-19 (Lucchese and Floel, 2020b). Both investigations identified potentially cross-reactive immune epitopes of human heat shock protein 90-beta (HSP90AB1) and SARS-CoV-2 proteins (Lucchese and Floel, 2020b; Marino Gammazza et al., 2020), and found that shared peptide sequences between heat shock protein 90-beta and SARS-CoV-2 nucleoprotein overlap with experimentally validated IEDB epitopes. Our analysis also identified potentially cross-reactive T-cell epitopes of the SARS-CoV-2 ORF6 protein and heat shock protein 90 proteins (HSP90AB1, HSP90AB3P, HSP90AA1, and HSP90AB2P) with predicted IC50 < 250nM for both peptides for at least one MHC allele. The predicted cross-reactive T-cell epitopes of the ORF6 protein and heat shock protein 90 proteins were overlapped with experimentally validated T-cell epitopes of SARS-CoV-1 or SARS-CoV-2 ORF6 proteins. Given the ubiquitous expression of heat shock protein 90 alpha (HSP90AA1) and beta (HSP90AB1) and increase in titers of autoantibody against HSP90AB1 in a small cohort of multisystem inflammatory syndrome in children (MIS-C) following SARS-CoV-2 infection (Gruber et al., 2020), it would be interesting to test the hypothesis that type IV hypersensitivity against heat shock protein 90 contributes to severe COVID-19 symptoms in the near future.

### Development of a web application for exploring potentially immunopathogenic SARS-CoV-2 epitopes and analyzing peptide vaccines against SARS-CoV-2

Due to a large number of predicted potentially cross-reactive B-cell and T-cell epitopes of SARS-CoV-2 and human proteins and the complexity of cross-reactivity prediction data, we developed an interactive web application for an interactive exploration of potential cross-reactivity of SARS-CoV-2 proteins or vaccines against SARS-CoV-2 and human proteins (**Fig. 3**). Our web application visualizes potentially cross-reactive B-cell and T-cell epitopes on the representative SARS-CoV-2 proteome alongside with known SARS-CoV-1/SARS-CoV-2 epitopes from Immune Epitope Database (IEDB) whose immunogenicity is experimentally validated. Additionally, user-provided peptide vaccine candidates against SARS-CoV-2 can be visualized and analyzed using the web application. By default, 820 peptide vaccine candidates against SARS-CoV-2 from OptiVax (Liu et al., 2020) is loaded.

**Figure 3.**
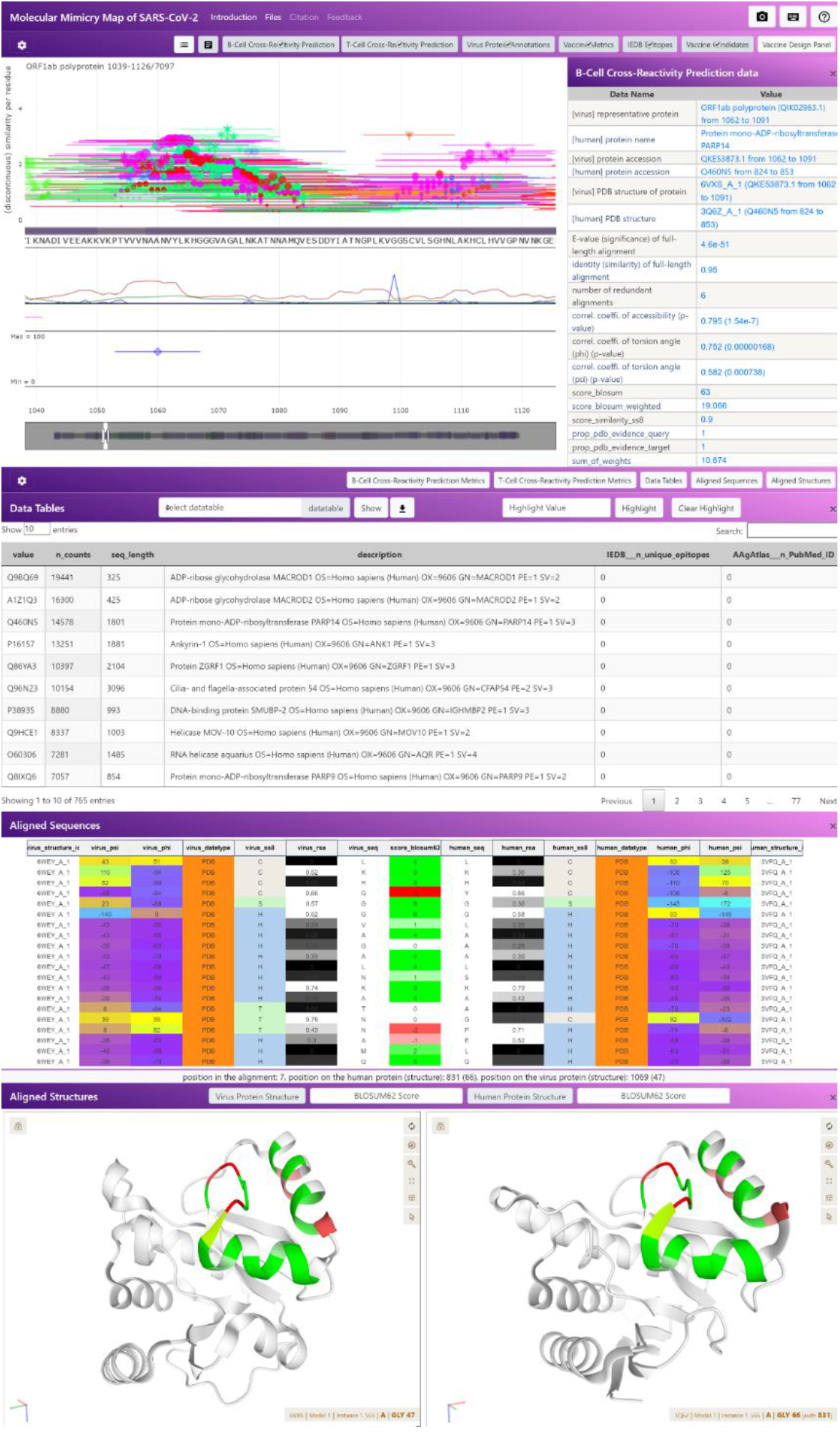
An overview of our interactive web application “Molecular Mimicry Map of SARS-CoV-2”. In the main panel (the graph at the top), a potentially cross-reactive B-cell epitope is represented by a line with a bubble in the middle of the line by default. The start and end of the line represent the start and end of the epitope on the representative SARS-CoV-2 protein. The color of the epitopes represents the name of a human protein from which the epitope was derived, and all B-cell or T-cell epitopes from the same human protein are visualized using the same color. Additional settings for the main panel can be found by clicking the setting icon in the top left corner of the main panel, where the window size for B-cell epitopes, number of T-cell and B-cell epitopes, and the size of the main graph can be adjusted. Visibility of additional tracks to the main graph, including relative surface availability (RSA) values of representative SARS-CoV-2 proteins, can be set in the main graph setting panel.

The main panel of our web application displays potentially cross-reactive B-cell and T-cell epitopes of SARS-CoV-2 proteins from 8,138 SARS-CoV-2 genomes. Because some SARS-CoV-2 proteins from 8,138 SARS-CoV-2 genomes are longer than its reference protein due to insertions, the longest protein belonging to each reference protein was chosen to represent the reference protein instead. Below the main graph panel, we added several additional panels for displaying supplementary information. Cross-reactive B-cell and T-cell epitopes displayed in the main graph can be filtered using the three parallel coordinate plots, in which users can set thresholds of several metrics to filter out less significant prediction results and adjust the sensitivity of cross-reactivity prediction during the analysis of peptide vaccine candidates. Additionally, potentially cross-reactive B-cell and T-cell epitopes can be filtered based on the gene expression patterns of human proteins in a selected set of tissues available in the Genotype-Tissue Expression (GTEx) dataset. When predicted cross-reactive epitopes visualized in the main graph is clicked, detailed information about the epitope is displayed in a table next to the main graph, and the aligned amino acid sequences of potentially cross-reactive epitopes of the SARS-CoV-2 and human proteins will be shown in the “Aligned Sequence” panel. When a B-cell epitope is clicked, if experimentally determined structures are available, the potentially cross-reactive B-cell epitopes will be displayed on 3D protein structures whose coordinates are automatically downloaded from either EMBL-EBI’s Coordinate Server (RCSB PDB database) or our GitHub repository (SARS-CoV-2 virus protein structure modeling results). Each row in the “Aligned Sequence” table represents a pair of matched amino acid residues of aligned SARS-CoV-2 and human epitopes, and the locations of the matched amino acids can be identified on the 3D protein structures by clicking the row, which will highlight the matched amino acid residue in either of 3D protein structures of SARS-CoV-2 and human proteins.

Currently, there are two licensed vaccines against SARS-CoV-2, one currently in use in the Chinese military and the other developed in Russia and licensed without completion of the Phase III trial (Krammer, 2020). Both vaccines are adenovirus-based non-replicating vector vaccines. Fourteen and nine vaccine candidates are currently undergoing Phase II and Phase III trials. Major hurdles of vaccine development against COVID-19 include antibody-dependent enhancement (ADE) and eosinophil-associated T_H_2-type immunopathology following exposure to SARS-CoV-2 after vaccination (Krammer, 2020; Lee et al., 2020; Lindsley et al., 2020). Recently, the Phase III trial of a leading vaccine candidate developed by the University of Oxford and AstraZeneca has been halted for six days in the United Kingdom after one participant experienced an adverse reaction during the trial (Mallapaty and Ledford, 2020; Phillips et al., 2020). The participant has developed symptoms of transverse myelitis and was later diagnosed with multiple sclerosis (MS), an autoimmune disease (Mallapaty and Ledford, 2020). An independent panel decided that the adverse reaction is unrelated to the vaccine trial, and the trial was resumed shortly in the UK, and a month later in the US. It has not been disclosed whether the participant received the vaccine or the placebo. Transverse myelitis is commonly associated with an autoimmune disease and bacterial and viral infections (Rodriguez et al., 2018), and rarely associated with vaccinations with hepatitis B virus (HBV), measles-mumps-rubella (MMR), and diphtheria-tetanus-pertussis (DTP) vaccines (Agmon-Levin et al., 2009). These symptoms could be involved with molecular mimicry and therefore vaccine candidates have to be carefully selected considering the possible cross-reactivity between SARS-CoV-2 and human proteins.

The web application enables researchers to explore potentially cross-reactive SARS-CoV-2 epitopes alongside custom peptide vaccines designed from the SARS-CoV-2 reference proteome. Our web application can be utilized to identify potentially suboptimal peptide vaccine candidates or less ideal part of a whole virus vaccine to design a safer vaccine for people with genetic and environmental predispositions to autoimmune diseases. The web application reports the number of overlapping potentially cross-reactive T-cell or B-cell epitopes for user-provided peptide vaccine candidates, which can inform the researchers for potential cross-reactivity of the peptide. Additionally, researchers can analyze vaccine candidates using the web application without privacy concern, since our web application run locally in the client’s web browser and was not designed to upload any data to external servers.

## Discussion

Antibody-dependent enhancement (ADE) describes an increased susceptibility to severe viral infection, paradoxically caused by the presence of neutralizing antibodies from previous exposure to or vaccination against the virus. ADE has been associated with flaviviruses, which include Dengue and Zika virus, and coronavirus. Dengue vaccination can increase the risk of dengue hospitalization (Hadinegoro et al., 2015), and an intermediate titer of neutralizing antibodies against the dengue virus has been reported to increase susceptibility to ADE (Katzelnick et al., 2017). The presence of sub-optimal antibodies against a virus or intermediate titer of neutralizing antibodies can increase the entry of the virus into cells expressing the Fc receptor, such as B cells, natural killer (NK) cells, macrophages, and neutrophils, exacerbating the disease. It has been reported that SARS-CoV-1 (Liu et al., 2019; Luo et al., 2018a; Yip et al., 2014) and MERS-CoV (Du et al., 2014; Wan et al., 2020) virus particles can enter a macrophage and affect its biological functions via its Fc receptor in the presence of antibodies against the virus.

During the development of a vaccine against SARS-CoV-1 and MERS, multiple studies reported increased infiltration of eosinophils in the lung or liver after challenged with the virus after vaccination despite the presence of neutralizing antibodies against the virus, leading to the development of hepatitis and pulmonary eosinophilia (PE) in animal models (Bolles et al., 2011; Czub et al., 2005; Deming et al., 2006; Tseng et al., 2012). The infiltration of eosinophils into the lung and exacerbated pneumonia were subsequently associated with the vaccination of nucleocapsid (N) protein of SARS-CoV-1 in two independent studies (Deming et al., 2006; Yasui et al., 2008). Interestingly, passive transfer of sera containing antibodies against the nucleocapsid protein (anti-N sera) into naïve mice did not reproduce the pulmonary eosinophilic infiltrates observed in mouse vaccinated with the nucleocapsid proteins (Deming et al., 2006), which led authors to conclude that the observed immunopathology with eosinophilic infiltrates was resulted from cell-mediated immune responses rather than from ADE, possibly involving type IV hypersensitivity reaction.

Dysregulated immune responses during SARS-CoV-2 infections have been associated with severe COVID-19 cases (Giamarellos-Bourboulis et al., 2020). Specifically, an elevated level of IL-6 (Chen et al., 2020) and decreased number of CD4+ and CD8+ T lymphocytes (lymphopenia) (Zhao et al., 2020) have been associated with severe COVID-19 cases. Also, there has been an increasing number of reports of autoimmune and inflammatory diseases following SARS-CoV-2 infections (Galeotti and Bayry, 2020), including multisystem inflammatory syndrome in children (MIS-C) (Akca et al., 2020; Gruber et al., 2020). Cross-reactivity against shared immune epitopes of proteins of human and invading pathogens, or molecular mimicry, is one of the suggested mechanisms that could initiate a series of dysfunctional immune responses that lead to the development of autoimmune diseases (Rojas et al., 2018). Therefore, we searched 3,323 non-redundant SARS-CoV-2 proteins from 8,138 SARS-CoV-2 genomes for possible immune epitope similarity with 20,595 human proteins.

Our findings demonstrate that the Macrodomain of nsp3 shares highly similar conformational epitopes (B-cell epitopes) and possibly cross-reactive MHC-I and MHC-II ligands (T-cell epitopes) with human PARPs (PARP14 and PARP9) and other human Macro domain proteins (MACROD1 and MACROD2). It has been reported that coronavirus infection can increase the expression of PARP enzymes in certain cell types *in vitro.* Grunewald reported that SARS-CoV-1 infection induces expressions of PARP9 and PARP14 in mouse brain CD11b+ cells (Grunewald et al., 2019), which is a marker for macrophage and microglial cells. Also, Heer and colleagues recently reported that SARS-CoV-2 infection-induced expression of PARP9 and PARP14 in normal human bronchial epithelial (NHBE) cells (Heer et al., 2020). PARP9 and PARP14 have regulatory roles in macrophage activation via STAT1 ADP-ribosylation (Iwata et al., 2016). According to the GTEx dataset, expressions of PARP9 and PARP14 are highest in lung, EBV-transformed lymphocytes, and whole blood tissues (www.gtexportal.org). Considering that autoantibodies against PARPs have been identified in SLE patients (Jeoung et al., 2004), it is tempting to speculate that the molecular mimicry between SARS-CoV-2 nsp3 and human PARP14 proteins might play a role in promoting inflammations and creating an inflammatory environment in the lung where PARP9 and PARP14 expressions are highest, which can lead to a series of dysfunctional immune response and eventually the development of autoimmune diseases in rare cases. Macrophage activation syndrome (MAS) has been suggested for its possible contribution to cytokine release syndrome in severe COVID-19 cases (Otsuka and Seino, 2020), which makes this hypothesis more tempting.

Other human proteins that share a significant number of cross-reactive T-cell and B-cell epitopes with SARS-CoV-2 nsp3 include MACROD1 and MACROD2. MARCOD1 is a nuclear-encoded mitochondrial protein, and it is mainly expressed in skeletal muscle, thyroid, and cardiac muscle tissues. There has been a report of a COVID-19 associated myositis case with severe muscle weakness (Zhang et al., 2020a). The patient has muscle weakness in her facial, bulbar, and proximal limb muscles with normal motor nerve conduction, and has autoantibodies such as anti-Ro/SSA (Sjögrensyndrome-related antigen) and anti-SAE 1 (Zhang et al., 2020a). A recent study suggested a possibly high frequency of paraspinal myositis and intramuscular edema (7 of 9 patients) in a small cohort of COVID-19 patients (n=9) (Mehan et al., 2020). Our finding that the Macro domain of MACROD1 and SARS-CoV-2 nsp3 proteins share a significant number of predicted cross-reactive immune epitopes might imply that cross-reactivity against human MACROD1 proteins after SARS-CoV-2 infection is responsible for the development of COVID-19 associated myositis.

A proteomic study of SARS-CoV-1 viral particles has revealed the presence of SARS-CoV-1 nsp3 proteins in its virion, and suggested nsp3’ role in SARS-CoV-1 virion assembly process (Neuman et al., 2008). The study found that SARS-CoV-1 virus particles (virions) are composed of structural proteins including nucleocapsid (N), membrane (M), and spike (S) proteins, as well as several nonstructural and accessory proteins including nsp3, nsp2, nsp5, ORF3a, and ORF9b. Also, a small amount of host membrane proteins was detected, estimated to be ~5% of the total protein of virion (Neuman et al., 2008). The study estimated that the amount of SARS-CoV-1 nsp3 protein in the virion is comparable to that of structural proteins, larger than the amount of nsp2, nsp5, ORF3a, ORF9b, and host membrane proteins in the virion. The authors have attributed the abundance of SARS-CoV-1 nsp3 in the virion to the ability of nsp3 to bind single-strand nucleic acids, including ssRNA, and the presence of a transmembrane domain in nsp3 (Neuman et al., 2008). The authors also explained the presence of ORF3a and ORF9b in the virion by their associations with host membranes. There are currently three inactivated whole virus particle vaccines against SARS-CoV-2 in Phase III clinical trials, four in Phase I/II clinical trials, and nine in its preclinical stages (Krammer, 2020). Also, there are one and three live attenuated SARS-CoV-2 vaccines under Phase I stage and preclinical stages, respectively (Krammer, 2020). Unlike spike-protein-based vaccines or its receptor-binding domain (RBD)-based vaccines, inactivated or live-attenuated SARS-CoV-2 vaccines are based on intact SARS-CoV-2 virions containing not only spike protein, but also other viral proteins including nsp3, N, M, ORF3a, and ORF9b proteins (Neuman et al., 2008). Moreover, the live attenuated vaccines (LAV) are capable of expressing nsp3 proteins in its infected cells.

Therefore, it would be better to significantly modify or remove the part of the SARS-CoV-2 genome encoding the Macro domain of nsp3 (ORF1 ab polyprotein from 1020 to 1197) to develop a vaccine that is potentially safer for a wider range of people, including people who are genetically and environmentally predisposed to autoimmune diseases. SARS-CoV-2 nonstructural protein 3 (nsp3) was suggested as a potential target for the development of vaccines against SARS-CoV-2 due to its high predicted protective antigenicity (Ong et al., 2020). SARS-CoV-2 nonstructural protein 3 is not well conserved in other coronaviruses that cause mild symptoms in human, but it is well conserved in coronaviruses that cause fatal infections, such as SARS-CoV-1 and MERS-CoV, and capable of binding poly(ADP-ribose) like other coronavirus nsp3 proteins via its Macrodomain (Frick et al., 2020; Ong et al., 2020). Nsp3 proteins in coronaviruses are capable of binding and hydrolyzing ADP-ribose from proteins, and it has been suggested that the enzymatic activity nsp3 is required for suppressing the innate immune response against viral infection mediated by PARPs (Grunewald et al., 2019). Inactivating mutations of the catalytic triad of nsp3 reduce the virulence of Mouse hepatitis virus strain JHMV (MHV), the prototypical coronavirus strain used in coronavirus studies, in bone marrow-derived macrophages (Grunewald et al., 2019). Grunewald and colleagues showed that knockdown of PARP14 or PARP12 increased proliferation of nsp3 mutant viruses in macrophages, but not wild-type coronavirus (MHV), implying the role of nsp3 in inhibition of PARP-mediated innate immune response (Grunewald et al., 2019). The nsp3 mutant viruses replicate poorly in the macrophages, but the mutations do not completely ablate the replication capacity of the virus. Therefore, the removal of the Macro domain of SARS-CoV-2 nsp3 in the SARS-CoV-2 genome seems feasible in the development of live-attenuated or inactivated SARS-CoV-2 vaccines.

Overall, we present a novel computational pipeline for accurate prediction of potentially cross-reactive conformational epitopes shared between human proteins and proteins of invading pathogen. To facilitate the development of a safe and effective vaccine against SARS-CoV-2, we applied our pipeline to predict potentially cross-reactive conformational epitopes and MHC ligands of SARS-CoV-2 and human proteomes. Additionally, we developed an interactive web application to visualize published and user-provided peptide vaccines against SARS-CoV-2 alongside the predicted cross-reactive immune epitopes of SARS-CoV-2 and human proteins. Our interactive web application can provide a foundation for the investigation of molecular mimicry in the pathogenesis of autoimmune disease following COVID-19.

## Methods

### Clustering and Alignment of 8,138 SARS-CoV-2 Proteomes

93,413 protein sequences derived from 8,138 sequenced SARS-CoV-2 (virus taxid:2697049, host taxid:9606) genomes were downloaded from NCBI Virus (www.ncbi.nlm.nih.gov/labs/virus/vssi/) (accessed 26 June 2020). The protein sequences were clustered with MMseqs2 easy-cluster (Steinegger and Soding, 2017) (float_thres_seq_identity = 0.4, float_thres_align_coverage = 0.8). resulting in 144 clusters.

Most clusters contained less than 10 proteins. Twelve clusters containing more than 7,000 proteins were selected for further analysis. These twelve clusters corresponded to twelve SARS-CoV-2 proteins: ORF1a polyprotein, ORF1ab polyprotein, surface glycoprotein (S), ORF3a protein, envelope protein (E), membrane glycoprotein (M), ORF6 protein, ORF7a protein, ORF7b protein, ORF8 protein, nucleocapsid phosphoprotein (N), and ORF10 protein. The longest protein sequence in each cluster was selected to represent the cluster. GenBank accessions of twelve representative protein sequences are the following: QJR92312.1, QIK02963.1, QJD23741.1, QJR84790.1, QJD24727.1, QKV08194.1, QKG86850.1, QJD47723.1, QKV36677.1, QKE12265.1, QJX68546.1, and BCF79872.1.

Proteins in each cluster were then aligned to the representative protein sequence by iteratively searching the proteins with a profile hidden Markov model of the representative sequence using the jackhmmer module (maximum four iterations) from HMMER 3.3 (Eddy, 2011) (released 18 November 2019). All 83,323 proteins in the twelve clusters were successfully aligned to representative sequence of each cluster.

### Sequence similarity search using BLAST and HMMER

A collection of human protein sequences representing a human proteome was downloaded from UniProt (proteome ID: UP000005640) (accessed 15 April 2020). A BLAST database of the 20,595 human proteins was made using the makeblastdb module, and the 93,413 SARS-CoV-2 proteins were searched against the database to find potentially homologous regions between SARS-CoV-2 proteins and human proteins (BLAST version 2.10.0+, released 16 December 2019).

A database of profile hidden Markov models (profile HMM) for every human protein sequence was built by iteratively searching a profile HMM for each human protein against all 2,949,581 protein sequences of viruses infecting human, downloaded from NCBI Virus (accessed 26 June 2020), using the jackhmmer module in HMMER 3.3 (maximum number of iterations = 5, E-value threshold = 30).

### Prediction of relative accessible surface area (RSA) and secondary structure of SARS-CoV-2 and human proteins using ACCpro 5 and SSpro 5

ACCpro 5.2 from SCRATCH-1D 1.2 suite (download.igb.uci.edu/) (Magnan and Baldi, 2014) (released November 2018) predicts relative accessible surface area (RSA) of every residue in a given protein sequence, respectively, through homology detection and neural networks. ACCpro 5.2 first uses PSI-BLAST (Position-Specific Iterative BLAST) and UniRef50 database (Suzek et al., 2015) to identify conserved domains on the given protein sequence, which is given as inputs to an ensemble of bidirectional recursive neural networks (BRNNs) to make an initial prediction of relative accessible surface area of the protein sequence. Next, ACCpro 5.2 combines the initial prediction with the actual relative accessible surface area of existing protein structures in the Protein Data Bank (PDB) database to make a final prediction of RSA. SSpro 5.2 from the same suite predicts secondary structure assignment of every residue in a given protein sequence using the same approach as ACCpro 5.2 through homology detection and an ensemble of neural networks.

Relative accessible surface area and secondary structure of 20,595 human protein sequences representing the human proteome were predicted using ACCpro/SSpro 5.2. ACCpro/SSpro 5.2 successfully predicted RSA and secondary structure of 20,593 human proteins but failed to predict RSA of the two largest human proteins, mucin-16 (14,507 amino acids) and titin (34,350 amino acids).

Relative accessible surface area and secondary structure of 83,323 clustered SARS-CoV-2 protein sequences were also predicted using ACCpro/SSpro 5.2. To reduce computation time, redundant protein sequences were dropped from the 83,228 SARS-CoV-2 protein sequences, resulting in 3,323 non-redundant SARS-CoV-2 protein sequences, comprising 969 ORF1a polyproteins, 1,336 ORF1ab polyproteins, 310 spike glycoproteins, and 708 other proteins (**Table S1**). SARS-CoV-2 ORF1a and ORF1ab genes encode two polyproteins, PP1a and PP1ab. SARS-CoV-2 PP1a and PP1ab are cleaved into 11 and 15 nonstructural proteins by proteolytic activities of two nonstructural proteins (nsp3 and nsp5), which are part of the two polyproteins. The redundancy of SARS-CoV-2 protein sequences was further reduced by cleaving ORF1a and ORF1ab polyproteins into 11 and 15 nonstructural proteins *in silico* and removing redundant nonstructural protein sequences. The rationale behinds this approach was the assumption that the tertiary structure of each nonstructural protein is largely independent of each other, and an amino acid variation in one nonstructural protein does not affect the tertiary structures of other nonstructural proteins. For *in silico* cleavage of ORF1a and ORF1ab polyproteins, 20 amino acids long sequences before the fifteen cleavage sites were retrieved from the UniProtKB COVID-19 resource (covid-19.uniprot.org) using the accessions P0DTC1 and P0DTD1 for ORF1a and ORF1ab, respectively. All 28,394 cleavage sites of non-redundant ORF1a and ORF1ab polyproteins were successfully identified by searching the polypeptide sequences with the 20 amino acids long sequences while allowing three mismatches and one deletion or insertion, identifying 30,699 nonstructural proteins. Removal of redundant nonstructural protein sequences resulted in 1,449 non-redundant SARS-CoV-2 nonstructural proteins (**Table S2**). ACCpro/SSpro 5.2 successfully predicted RSA and secondary structure of the 1,449 non-redundant SARS-CoV-2 nonstructural proteins and 1,018 non-redundant SARS-CoV-2 proteins other than ORF1a and ORF1ab polyproteins.

### Relative accessible surface area (RSA) calculation and secondary structure assignment of RCSB Protein Data Bank (PDB) database

Research Collaboratory for Structural Bioinformatics Protein Data Bank (RCSB PDB) curates and annotates experimentally determined protein structures, including 291 recently determined SARS-CoV-2 protein structures. The Worldwide PDB (wwPDB) (www.wwpdb.org/) maintains the PDB archive and provides access to the database via its FTP server (ftp.wwpdb.org). The protein structures in the PDB archive are updated every Wednesday and are provided in three formats: XML, PDB, and mmCIF format. The PDB file format is created in 1970 to represent macromolecular structure data and is currently most widely used to represent protein structures. However, due to the fixed column length of the PDB format, a protein structure with more than 9,999 atoms or a chain identifier longer than two letters cannot be stored in the format. Consequently, structure data of hundreds of protein structures, including a couple of SARS-CoV-2 protein structures like 6WKS (www.rcsb.org/structure/6WKS), is not available in the PDB format. In contrast, the column lengths in the mmCIF format are not fixed, allowing a mmCIF file to store a structure composed of an unlimited number of atoms. Therefore, total 166,891 macromolecular structures in the mmCIF format were downloaded from the PDB archive (accessed 10 July 2020) using the script provided by the RCSB PDB ftp download page (www.rcsb.org/pages/download/ftp).

First, 166,891 macromolecular structures in the mmCIF format were loaded into a Python environment (Python 3.7.6). Specifically, tabular data in the ‘atom_site’ data category was retrieved from each mmCIF file. Next, the macromolecular structures were cleaned by removing atoms that do not constitute a protein. The 3D coordinates of atoms that do not belong to one of the twenty standard amino acids were discarded, including atoms of cofactors and ligands bound to proteins and carbohydrates linked to glycosylated residues. Also, 3D coordinates of atoms belonging to a standard amino acid bound to proteins as a ligand and thus not a part of a protein were discarded. Additionally, when alternative 3D coordinates are available for an amino acid residue, only the first set of 3D coordinates was retained, and all other alternative sets of 3D coordinates were discarded.

In addition to the cleaning, a couple of data attributes were modified to make annotations of protein structures more consistent across the PDB archive. The first modified attribute is ‘auth_seq_id’, which contains residue numbers provided by authors who published the structure. The residue numbers in the ‘auth_seq_id’ attribute are sometimes inconsistent with the order of residues in a protein or not unique to a residue in a protein, often when the region of protein does not normally present in the original protein of interest, such as residues belonging to a His-tag or an inserted amino acid sequence that do not present in the original protein. However, residue numbers in the ‘label_seq_id’ attribute are always consistent with the order of residue in a protein and unique to each residue. One of the downstream software tools, mkdssp, uses residue numbers in the ‘auth_seq_id’ attribute to assign residue numbers to its secondary structure classification result. Therefore, residue numbers in the ‘auth_seq_id’ attribute were replaced with the residue numbers in the ‘label_seq_id’ attribute.

The second modified attribute is ‘pdbx_PDB_model_num’, which contains model numbers for multiple protein models present in a macromolecular structure. Protein structures determined by nuclear magnetic resonance (NMR) spectroscopy often contain multiple models representing possible protein structures satisfying the conformational constraints observed in the NMR experiment. In contrast, protein structures determined by X-ray crystallography contain a single protein model, and the ‘pdbx_PDB_model_num’ attribute is often omitted. For protein structures lacking ‘pdbx_PDB_model_num’ attribute, the attribute was added to the structure and filled with 1, representing the first model.

After cleaning and preprocessing of mmCIF files were completed, each polypeptide in a mmCIF file is written as an individual mmCIF file. One of the downstream analysis tools, mkdssp discards all atoms that do not belong to the first model (ignore atoms with ‘pdbx_PDB_model_num’ attribute other than 1) when calculating the structural properties of a protein. Therefore, each mmCIF file containing multiple models (mostly structures determined by the NMR experiment) was split into multiple mmCIF files for each model. Additionally, every ‘pdbx_PDB_model_num’ attribute of split models is replaced with 1, allowing mkdssp program to calculate the structural property of every model in a protein structure. In total 769,761 unique structures of polypeptides were retrieved and written as an equal number of separate mmCIF files.

Next, structures with missing atoms were detected by counting the number of atoms for each residue. 167,771 polypeptide structures contained residues with missing atoms. Missing atoms in these structures were filled with complete_pdb function in a MODELLER python package (version 9.24, released 6 April 2020) (Webb and Sali, 2016). Among 167,771 polypeptide structures with missing atoms, 167,758 structures were successfully corrected with MODELLER.

After correcting residues with missing atoms, the mkdssp program was used to calculate the structural properties of 769,748 polypeptide structures. Structural properties of 769,676 polypeptide structures were successfully calculated by mkdssp, and the output files are loaded into the Python environment for further analysis. mkdssp calculates solvent accessible surface area (ASA) of each residue. The ASA values in the mkdssp output were divided by maximum allowed solvent accessibility (MaxASA) values empirically determined by Tien et. al. (Tien et al., 2013) to calculate relative solvent accessibility (RSA) for each residue. mkdssp also calculates torsion angles phi and psi for each residue. mkdssp then use the calculated torsion angles to assign one of the eight types of secondary structure to each residue: helix (G), α-helix (H), π-helix (I), β-stand (E), bridge (B), turn (T), bend (S), and undefined (C). Each sequencing quality score in FASTQ format is encoded in a single ASCII character using the Phred+33 encoding. Similarly, the calculated RSA, torsion angles phi and psi, and the assigned secondary structure type for each residue were encoded into one or two ASCII characters. For RSA and torsion angles, two ASCII characters were used to encode each value, while only one ASCII character was used to encode secondary structure type. Additionally, the amino acid sequence of a polypeptide structure was retrieved from the mkdssp output. In a protein structure determined by X-ray crystallography, 3D coordinates of atoms in the flexible loops are often not available, which creates a gap in the structure. In the amino acid sequence retrieved from the mkdssp output, such gaps were filled with ‘X’ so that BLASTP program can more accurately align the sequence from the protein structure to the original protein sequence. All the information extracted from the mkdssp output is saved to the file system using FASTA format.

### Relative accessible surface area (RSA) calculation and secondary structure assignment of SARS-CoV-2 proteins by template-based protein structure modeling

Through homology modeling, protein structures of SARS-CoV-2 proteome can be predicted by using homologous proteins with known structures as templates. Predicted protein structures of SARS-CoV-2 proteome were retrieved from three different sources: the SWISS-MODEL repository for SARS-CoV-2 proteome (Waterhouse et al., 2018) (swissmodel.expasy.org/repository/species/2697049) (accessed 15 July 2020), SARS-CoV-2 structure modeling results from C-I-TASSER pipeline (Huang et al., 2020) (zhanglab.ccmb.med.umich.edu/COVID-19/) (released 6 May 2020), and SARS-CoV-2 structure modeling results from RaptorX pipeline and refinement of Google’s AlphaFold SARS-CoV-2 protein structure models (Heo and Feig, 2020) (github.com/feiglab/sars-cov-2-proteins) (accessed 16 July 2020). In total, 140 predicted protein structures in PDB format were retrieved (73 from SWISS-MODEL, 43 from Feig_Lab, 24 from Zhang_Lab). These predicted protein structures were cleaned, modified, and split into individual polypeptide chains, as previously described for the processing of the PDB database. A total of 231 individual polypeptide structures were retrieved, which was subsequently processed by mkdssp to calculate structural properties of each polypeptide structure. As previously described, RSA, torsion angles phi and psi, and secondary structure type were retrieved from the mkdssp outputs and saved as FASTA format files.

### Calculation of consensus relative accessible surface area (RSA) and structural properties of SARS-CoV-2 and human proteins

To calculate consensus structural properties of human and SARS-CoV-2 virus proteins, protein sequences were searched against the database of protein sequences with predicted or known structures. 3,323 non-redundant SARS-CoV-2 protein sequences and 20,531 human protein sequences were searched against a database of 769,676 protein sequences with experimentally determined protein structures. Additionally, the non-redundant SARS-CoV-2 proteins were searched against a database of 231 protein sequences with predicted structures from homology modeling.

Overall, 2,052 and 408,183 PDB structures were aligned against SARS-CoV-2 and human protein sequences (e-value threshold 0.05). All 231 homology-modeled SARS-CoV-2 protein structures were aligned against SARS-CoV-2 virus proteins (e-value threshold 0.05). Consensus structural properties were then assigned to each residue by using structural properties of aligned structures. Weighted averages of relative solvent accessibility and torsion angles ϕ (phi) and ψ (psi) of aligned structures were calculated to retrieve consensus structural properties. To retrieve consensus secondary structure classifications, a secondary structure classification with the highest sum of weights was retrieved for each position.

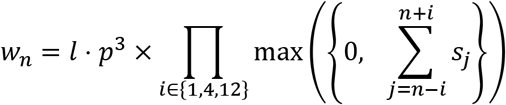

*w_n_*: weight at the n-th position of an alignment between structure and protein sequence

*s_i_*: BLOSUM62 score at the jth position of the alignment

*l:* length of the alignment

*p:* the identity of the alignment (number of identical residues/length of the alignment)

### Prediction of cross-reactive MHC class I and class II epitopes of SARS-CoV-2 and human proteins

224,400 non-redundant pairs of SARS-CoV-2 and human peptide sequences were extracted from the 150,744 alignments between human and SARS-CoV-2 protein sequences using the window sizes of 8, 9, 10, and 15 amino acids. To predict the cross-reactive T-cell epitopes in a reasonable timeframe, only the MHC alleles that have been linked to autoimmune diseases were used for the prediction. Using the list summarized by Vasiliki Matzaraki (Matzaraki et al., 2017), a list of 34 MHC alleles (18 MHC Class I alleles, 16 MHC Class II alleles) was retrieved. The MHC alleles in the list have been linked to 10 autoimmune diseases in total, which are following: rheumatoid arthritis, celiac disease, psoriasis, ankylosing spondylitis, systemic lupus erythematosus, type 1 diabetes, multiple sclerosis, Crohn’s disease, ulcerative colitis, and Graves’ disease (**Table S3**). Also, only SARS-CoV-2 and human peptide pairs with significant similarity scores (about 20,000 pairs for both MHC classes) were subjected to MHC-binding predictions to further reduce the computation time.

pVACtools version 1.5.9 (Hundal et al., 2020) was installed via Docker. In order to make MHC binding prediction more robust, all twelve algorithms in pVACtools were utilized to predict a binding affinity using the consensus of different algorithms (Jurtz et al., 2017; Shao et al., 2020; Zhang et al., 2009). For MHC class I and class II binding predictions, eight and four different algorithms were utilized, respectively: MHCflurry, MHCnuggetsI, NetMHC, NetMHCcons, NetMHCpan, PickPocket, SMM, and SMMPMBEC for MHC class I binding prediction and MHCnuggetsII, NetMHCIIpan, NNalign, and SMMalign for MHC class II binding prediction. A geometric mean of predicted affinity (IC50) values from different algorithms were calculated for each peptide to calculate the consensus of multiple MHC binding prediction algorithms. Overall, MHC binding affinities of 23,921, 27,230, 23,577, and 17,557 pairs of aligned human and virus peptides of 8, 9, 10, 15 amino acids in lengths were predicted for 34 autoimmune-associated MHC alleles using the consensus of a maximum of eight algorithms. The list of autoimmune-associated MHC class I alleles are the following: A*02:01, A*24:02, B*08:01, B*13:02, B*18:01, B*27:02, B*27:05, B*38:01, B*39:06, B*40:01, B*40:02, B*44:02, B*47:01, B*50:01, B*51:01, C*06:02, C*12:02, C*12:03. The list of autoimmune-associated MHC class II alleles are the following: DPB1*01:01, DPB1*04:02, DPB1*05:01, DQA1*01:02, DQA1*05:01, DQB1*02:01, DQB1*03:02, DQB1*06:02, DRB1*01:03, DRB1*03:01, DRB1*04:01, DRB1*04:04, DRB1*08:01, DRB1*13:03, DRB1*14:01, DRB1*15:01.

A potentially cross-reactive pair of T-cell epitopes derived from human and virus proteins should strongly bind an MHC molecule with high binding affinity. Therefore, the product of predicted MHC binding affinities (IC50) of human and virus peptides was used to identify a cross-reactive pair of T-cell epitopes. 48,497 potentially cross-reactive pairs of T-cell epitopes with a product of predicted binding affinity values IC50 (in nM) smaller than 250000 (nM^2^) were retrieved. The identified potential cross-reactive pairs of T-cell epitopes were mapped to the consensus SARS-CoV-2 protein sequences, which can be visualized in the web application.

### Alignment of experimentally validated IEDB epitopes to consensus SARS-CoV-2 proteome

Immune Epitope Database and Analysis Resource (IEDB) (www.iedb.org) curates experimentally validated B-cell and T-cell antigens from publications or submitted datasets (Vita et al., 2019). An entire IEDB database was downloaded as several tabular datasets (accessed 18 August 2020). To retrieve epitopes of SARS coronavirus (SARS-CoV) and Severe acute respiratory syndrome coronavirus 2 (SARS-CoV-2), epitopes with parent organism taxonomy identifier NCBI:txid694009 (Severe acute respiratory syndrome-related coronavirus) were selected for further analysis. Total 3,875 epitopes were retrieved, which were subsequently aligned against consensus SARS-CoV-2 proteome. As a result, 3,551 epitopes were successfully aligned to consensus SARS-CoV-2 proteome without gaps. Among these, 2,614 epitopes have higher than 80% similarity to the consensus protein sequence. 333 epitopes were derived from SARS-CoV-2, but the other 2,281 epitopes were derived from the other SARS-related coronaviruses, 1,512 of which were derived from SARS-CoV tor2 strain, a SARS-CoV strain isolated in Toronto, Canada. Since the linear sequences of these epitopes were significantly similar to the consensus protein sequences of SARS-CoV-2, and these epitopes were included in the subsequent analysis. 174 and 351 epitopes have positive B-cell and T-cell based assay results, respectively. Also, 1,558 epitopes have positive MHC-binding assay results. For each position in the consensus SARS-CoV-2 genome, a total number of positive B-cell and T-cell based assays of epitopes covering the position were counted, which can be visualized in the web application.

### Immunogenicity prediction of consensus SARS-CoV-2 proteome with broad coverage of the global population

From the multiple sequence alignment of ~8,000 SARS-CoV-2 proteomes, consensus SARS-CoV-2 proteome sequence was retrieved. Also, the number of non-consensus amino acid residues or gaps in the multiple sequence alignment was counted for each position of consensus protein sequences, which can be visualized in the web application. The vaccine designing process should be aware of these positions since vaccine harboring variable residues is less likely to elicit the same immune response to a broad range of SARS-CoV-2 viruses.

Using the consensus SARS-CoV-2 protein sequences, potentially immunogenic epitopes were predicted using an immunoinformatic approach. First, MHC class I and class II binding peptides were predicted using MHCflurry 2.0 (O’Donnell et al., 2020) and NetMHCIIpan-4.0 (Reynisson et al., 2020), respectively. Allele frequency data of MHC class I and class II alleles in various populations were retrieved from the immune epitope database (IEDB) population coverage tool (Bui et al., 2006), which itself retrieves its population information from Allele Frequency Net Database (AFND). Using the allele frequency data, the list of common MHC class I and II alleles were retrieved. For MHC class I alleles, there were 8, 100, 308, and 537 alleles with allele frequency greater than 0.5, 0.1, 0.01, or 0.001 in at least one of the populations available in the data. Potential MHC class I binding peptides of 8 - 12 amino acids were generated from the consensus SARS-CoV-2 protein sequences excluding gaps. For each allele frequency threshold value, the number of peptides with predicted binding affinity (IC50) < 250nM was counted for each position of SARS-CoV-2 consensus protein sequences, which can be visualized in the web application. MHCflurry also predicts MHC class I antigen processing scores in addition to binding affinity for an MHC class I molecule encoded by a specific allele. The model is trained with every eluted MHC I ligands identified by mass spectrometry. Thus, the antigen processing score is not specific to a specific MHC I molecules, but applicable to any MHC I allele. The average antigen processing scores for each position of SARS-CoV-2 consensus protein sequences were calculated, which can be visualized in the web application.

The peptide-binding groove of MHC class I molecules is located on the alpha chain, while that of MHC class II molecules is located between the alpha and beta chains. Therefore, peptide-binding characteristics of an MHC class II molecule are determined by two alleles encoding the alpha and beta chains. The data from the IEDB population coverage tool provided an unlinked allele frequency data separate for alpha and beta chains. Thus, for the prediction of peptides that have high binding affinity to common MHC class II molecules, the list of alleles encoding alpha and beta chains whose allele frequency is greater than 0.5, 0.1, 0.01, and 0.001 in at least one of the populations were retrieved. Next, all possible combinations of the common alpha and beta chain alleles retrieved from the earlier step were used to predict peptides that are likely to bind common MHC class II molecules. Peptides of 15 amino acids in length were generated from the consensus SARS-CoV-2 protein sequences, and the number of peptides with predicted binding affinity (IC50) < 250nm was counted for every position of SARS-CoV-2 consensus protein sequences, which can be visualized in the web application.

Next, potentially immunogenic B-cell linear epitopes of SARS-CoV-2 proteins were predicted using BepiPred 2.0 (Jespersen et al., 2017) server. The consensus SARS-CoV-2 protein sequences were given as an input file to the BepiPred 2.0 server. Since the server did not receive the protein longer than 6,000 amino acids, the consensus protein sequence encoded by the ORF1ab gene (7,097 amino acids in length) was split into two protein sequences containing nsp1-10 protein sequences and nsp11-16 protein sequences, respectively. The B-cell linear epitope prediction score was retrieved for each position on the consensus SARS-CoV-2 protein sequences, which can be visualized in the web application.

### Development of “3M of SARS-CoV-2” web application

In order to provide an interactive interface to explore the identified potential B-cell and T-cell cross-reactive epitope pairs from human and SARS-CoV-2 proteins, a web-based application was designed. The web application can be used to visualize the already designed peptide vaccine to filter out peptides that can potentially cause immunopathological reactions in genetically and environmentally susceptible individuals. The web application locally processes user-provided FASTA files containing designed peptide vaccines and provides an interface to analyze and filter the peptide vaccine candidates according to the number of overlapped cross-reactive regions and other metrics useful for peptide vaccine design, including the number of mutations at each position. Researchers can use the web application to visualize the protein structures of potentially crossreactive human and SARS-CoV-2 proteins, the alignment between human and virus proteins, and structural properties of the aligned sequences.

To visualize protein structures in the web application, PDBe Molstar (Mol*) JS plugin version 1.1.0 was used. Coordinates of a protein structure in the PDB database is downloaded from EMBL-EBI’s Coordinate Server (ebi.ac.uk/pdbe/coordinates/) and visualized in the web application. For drawing interactive scatter plots, parallel coordinate plots, and the scrollable table, Plotly.js version 1.54.7 was used. To draw a static table, D3.js version 3.5.5 was utilized. To parse tabular data files hosted on the web, PapaParse.js version 5.2.0 and Dataframe.js version 1.4.3 were used. For displaying interactive data tables, DataTables.js version 1.10.21 was used. For performing large numeric operations, NumJS version 0.15.1 was utilized. To create colormaps, Chroma-js version 2.1.0 was used. To save a local Javascript object in the web browser as a file, FileSaver.js version 1.3.4 was used. To capture the current view of the web application and save the screenshot as an image file, html2canvas.js version 1.0.0-rc.7 was used. Lastly, Bootstrap version 4.5.2 with JQuery version 3.5.1 was utilized to add components such as buttons and dialog boxes to the web application. Additionally, Google Material Design’s icons were utilized to add icons in some of the buttons in the web application.

## Supplementary Materials

**Figure S1.**
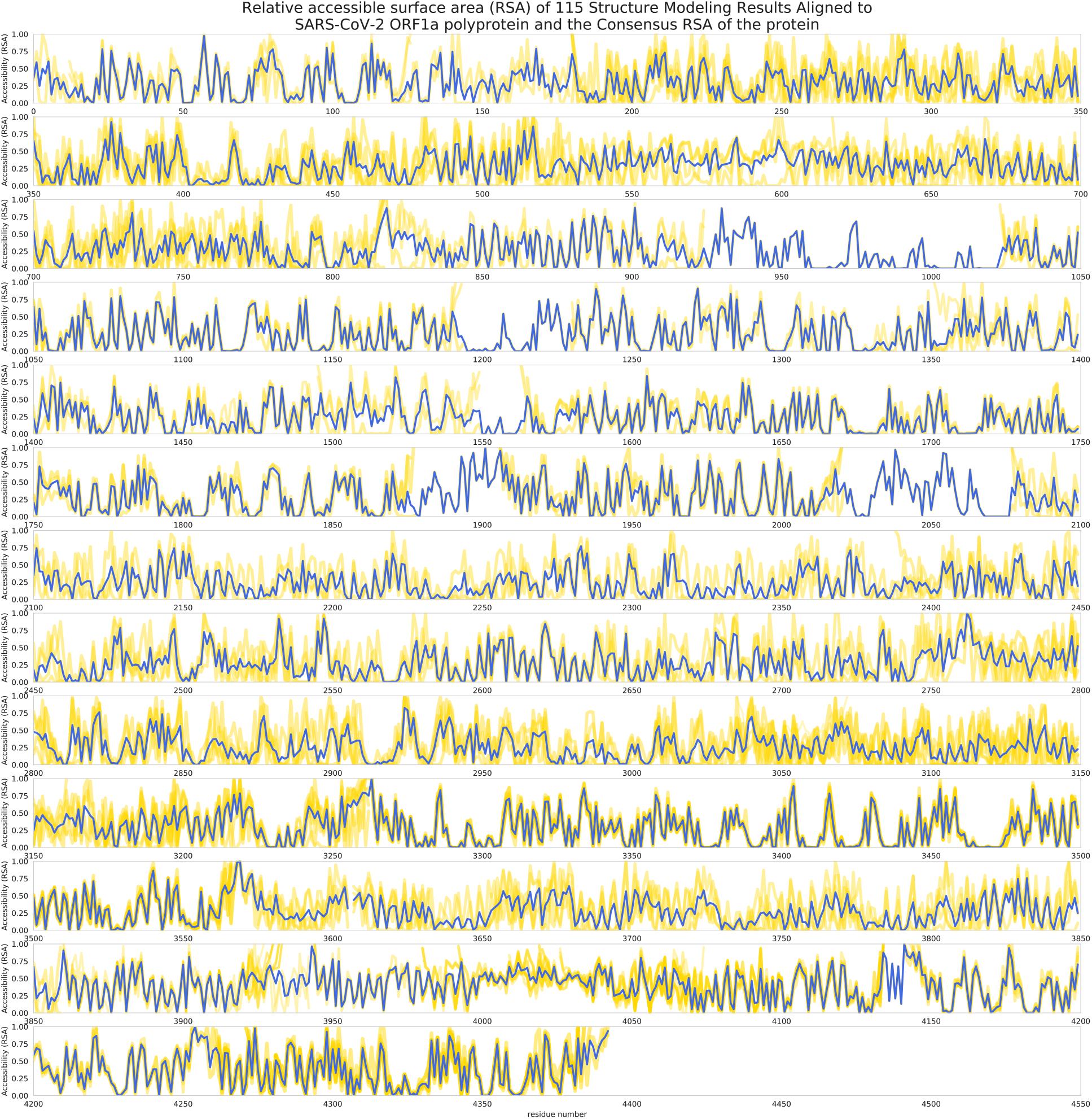
Consensus relative accessible surface area (RSA) of a representative SARS-CoV-2 ORF1a polyprotein. Consensus surface accessibilities of 3,323 non-redundant SARS-CoV-2 proteins were first calculated from available experimentally determined protein structures. For residues that were not covered by experimentally determined structures, high-quality protein structure modeling results of SARS-CoV-2 proteins were used instead. The yellow lines represent surface accessibilities of each protein structure, while the blue line represents the consensus surface accessibilities calculated for each residue.

**Figure S2.**
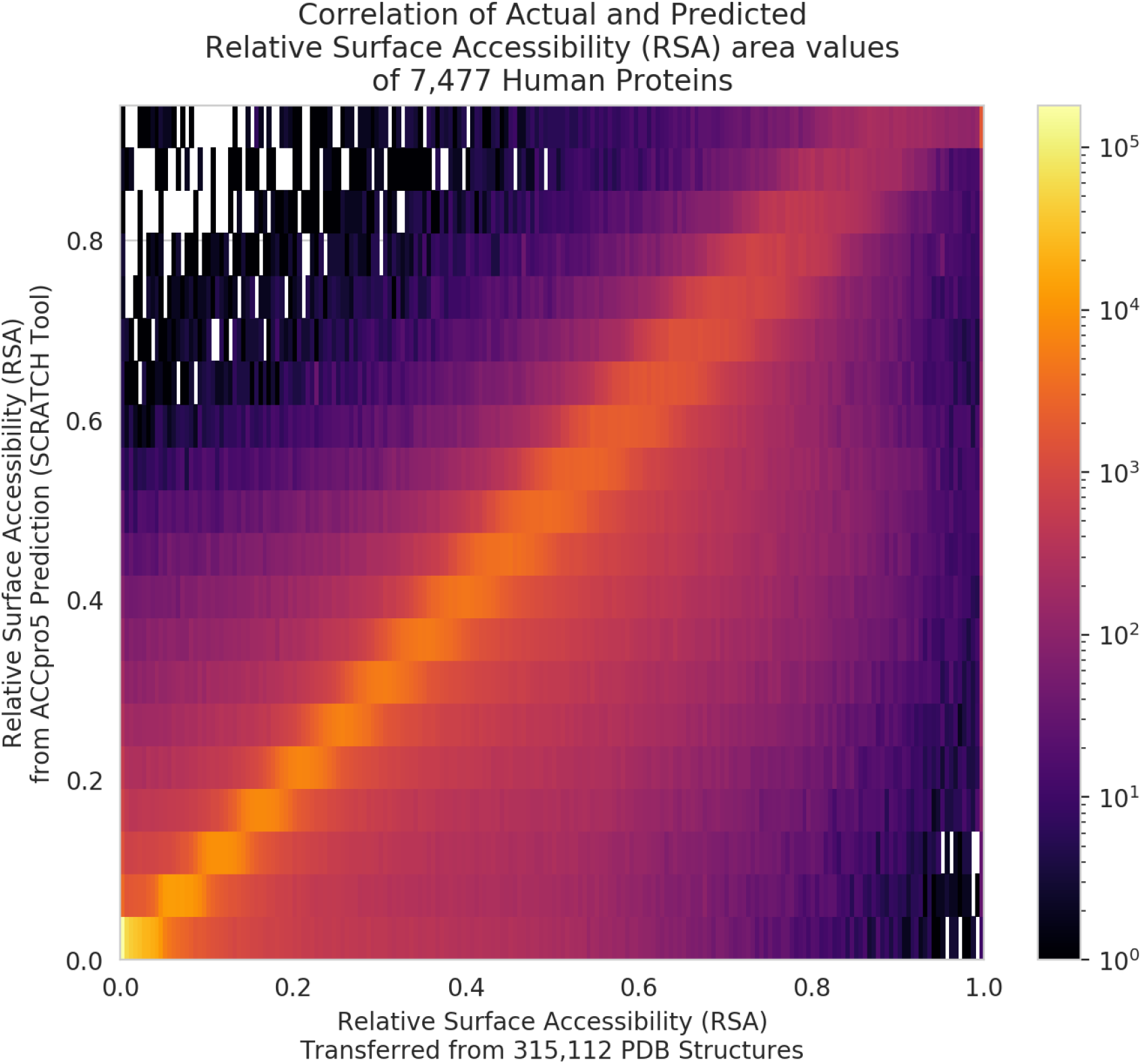
General agreements between ACCpro5-predicted relative surface accessibility (RSA) values and actual RSA values calculated from experimentally determined human protein structures. The heatmap represents a 2D histogram of all amino acid residues of human proteomes for which experimentally determined protein structures are available for calculation of actual RSA values, binned by predicted RSA values and actual RSA values for Y- and X-axis, respectively. Experimentally determined protein structures were available for 7,477 human proteins.

**Figure S3.**
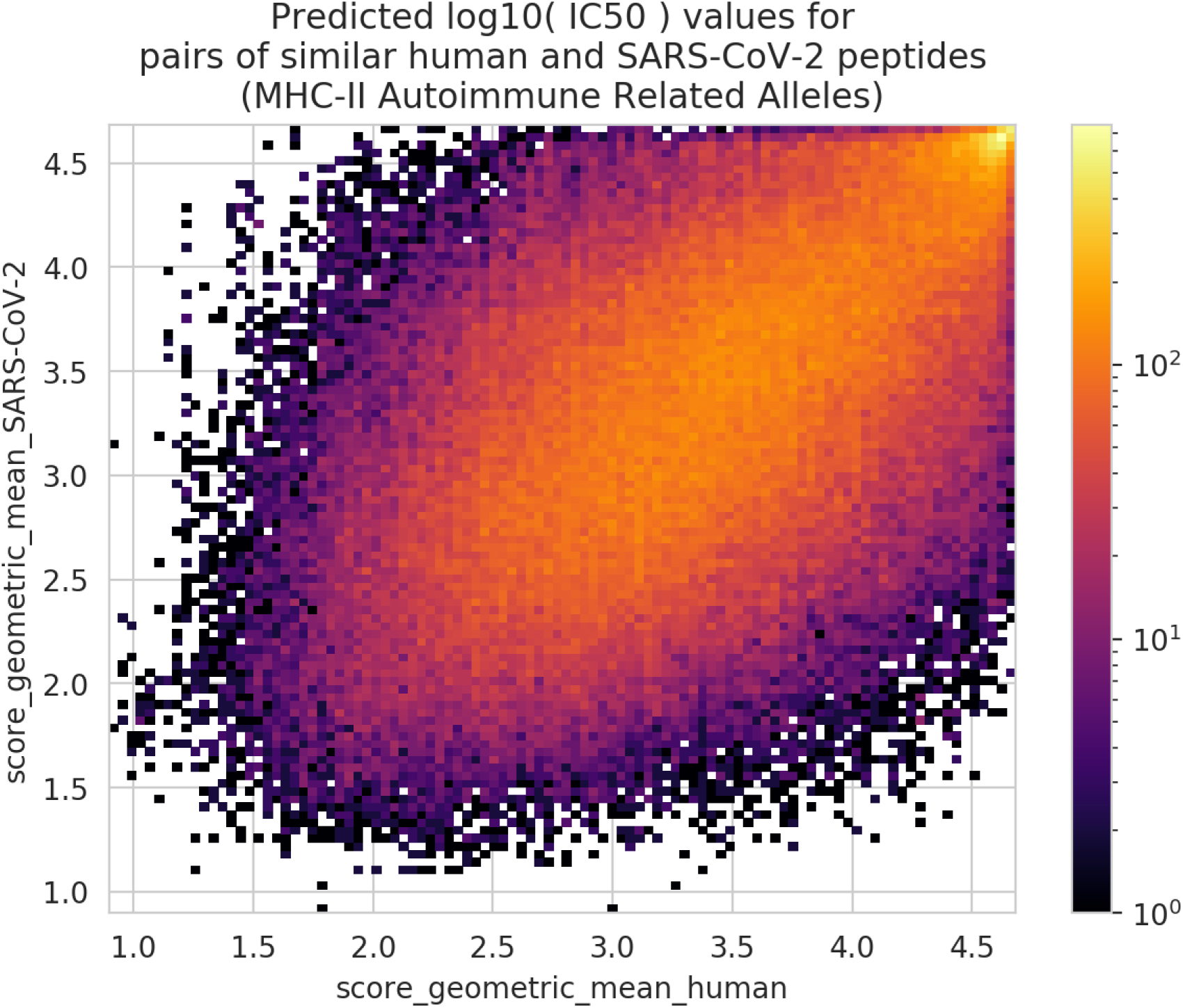
Predicted IC50 values of potentially cross-reactive CD4+ T-cell epitopes of human and SARS-CoV-2 proteins. We calculated an MHC-binding affinity of each peptide using geometric averages of IC50 values from up to eight distinct algorithms.

**Table S1.** Number of non-redundant SARS-CoV-2 proteins analyzed in this study.

| Protein name | Number of non-redundant proteins |
| --- | --- |
| ORF1a polyprotein | 969 |
| ORF1ab polyprotein | 1336 |
| surface glycoprotein (S) | 310 |
| ORF3a protein | 180 |
| envelope protein I | 29 |
| membrane glycoprotein (M) | 64 |
| ORF6 protein | 39 |
| ORF7a protein | 58 |
| ORF7b protein | 22 |
| ORF8 protein | 71 |
| nucleocapsid phosphoprotein (N) | 218 |
| ORF10 protein | 27 |

**Table S2.** Number of non-redundant SARS-CoV-2 non-structural proteins from ORF1a and ORF1ab polyproteins after proteolytic cleavage.

| Nonstructural protein name | Number of non-redundant proteins |
| --- | --- |
| nsp1 | 70 |
| nsp2 | 199 |
| nsp3 | 417 |
| nsp4 | 88 |
| nsp5 | 64 |
| nsp6 | 57 |
| nsp7 | 16 |
| nsp8 | 39 |
| nsp9 | 17 |
| nsp10 | 13 |
| nsp11 | 2 |
| nsp12 | 137 |
| nsp13 | 117 |
| nsp14 | 94 |
| nsp15 | 69 |
| nsp16 | 50 |

**Table S3.** List of major histocompatibility complex (MHC) alleles that has been associated with autoimmune disease by fine-mapping studies. This table is derived from a table compiled by Matzaraki and colleagues (Matzaraki et al., 2017).

| Disease Name | MHC Allele | MHC class | PubMed ID |
| --- | --- | --- | --- |
| rheumatoid arthritis | HLA-B*40:02 | I | 27486778 |
| celiac disease | HLA-B*08:01 | I | 25894500 |
| celiac disease | HLA-B*39:06 | I | 25894500 |
| psoriasis | HLA-C*06:02 | I | 25087609 |
| psoriasis | HLA-C*12:03 | I | 25087609 |
| ankylosing spondylitis | HLA-B*27:02 | I | 25994336 |
| ankylosing spondylitis | HLA-B*27:05 | I | 25994336 |
| ankylosing spondylitis | HLA-A*02:01 | I | 25994336 |
| ankylosing spondylitis | HLA-DRB1*01:03 | II | 25994336 |
| ankylosing spondylitis | HLA-B*13:02 | I | 25994336 |
| ankylosing spondylitis | HLA-B*40:01 | I | 25994336 |
| ankylosing spondylitis | HLA-B*40:02 | I | 25994336 |
| ankylosing spondylitis | HLA-B*47:01 | I | 25994336 |
| ankylosing spondylitis | HLA-B*51:01 | I | 25994336 |
| systemic lupus erythematosus | HLA-DQB1*06:02 | II | 25533202 |
| systemic lupus erythematosus | HLA-DRB1*15:01 | II | 25533202 |
| systemic lupus erythematosus | HLA-DRB1*03:01 | II | 17997607 |
| systemic lupus erythematosus | HLA-DRB1*03:01 | II | 23084292 |
| systemic lupus erythematosus | HLA-DRB1*08:01 | II | 23084292 |
| systemic lupus erythematosus | HLA-DQA1*01:02 | II | 23084292 |
| systemic lupus erythematosus | HLA-DRB1*15:01 | II | 26808113 |
| systemic lupus erythematosus | HLA-DQB1*06:02 | II | 26808113 |
| type 1 diabetes | HLA-B*39:06 | I | 26168013 |
| type 1 diabetes | HLA-B*18:01 | I | 26168013 |
| type 1 diabetes | HLA-B*50:01 | I | 26168013 |
| type 1 diabetes | HLA-DPB1*04:02 | II | 26168013 |
| type 1 diabetes | HLA-DPB1*01:01 | II | 26168013 |
| type 1 diabetes | HLA-A*24:02 | I | 26168013 |
| multiple sclerosis | HLA-DRB1*15:01 | II | 24278027 |
| multiple sclerosis | HLA-DRB1*03:01 | II | 24278027 |
| multiple sclerosis | HLA-DRB1*13:03 | II | 24278027 |
| multiple sclerosis | HLA-DRB1*04:04 | II | 24278027 |
| multiple sclerosis | HLA-DRB1*04:01 | II | 24278027 |
| multiple sclerosis | HLA-DRB1*14:01 | II | 24278027 |
| multiple sclerosis | HLA-A*02:01 | I | 24278027 |
| multiple sclerosis | HLA-DRB1*15:01 | II | 26343388 |
| multiple sclerosis | HLA-DRB1*03:01 | II | 26343388 |
| multiple sclerosis | HLA-DRB1*13:03 | II | 26343388 |
| multiple sclerosis | HLA-DRB1*08:01 | II | 26343388 |
| multiple sclerosis | HLA-DQB1*03:02 | II | 26343388 |
| multiple sclerosis | HLA-B*44:02 | I | 26343388 |
| multiple sclerosis | HLA-B*38:01 | I | 26343388 |
| Crohn's disease | HLA-DRB1*01:03 | II | 25559196 |
| Crohn's disease | HLA-C*06:02 | I | 25559196 |
| ulcerative colitis | HLA-DRB1*01:03 | II | 25559196 |
| ulcerative colitis | HLA-C*12:02 | I | 25559196 |
| Graves' disease | HLA-DRB1*03:01 | II | 26029868 |
| Graves' disease | HLA-DQA1*05:01 | II | 26029868 |
| Graves' disease | HLA-DQB1*02:01 | II | 26029868 |
| Graves' disease | HLA-DPB1*05:01 | II | 26029868 |

## References

Agmon-Levin, N., Kivity, S., Szyper-Kravitz, M., and Shoenfeld, Y. (2009). Transverse myelitis and vaccines: a multi-analysis. Lupus 18, 1198–1204.

Akca, U.K., Kesici, S., Ozsurekci, Y., Aykan, H.H., Batu, E.D., Atalay, E., Demir, S., Sag, E., Vuralli, D., Bayrakci, B., et al. (2020). Kawasaki-like disease in children with COVID-19. Rheumatol Int.

Albiol, N., Awol, R., and Martino, R. (2020). Autoimmune thrombotic thrombocytopenic purpura (TTP) associated with COVID-19. Ann Hematol 99, 1673–1674.

Amedei, A., Bergman, M.P., Appelmelk, B.J., Azzurri, A., Benagiano, M., Tamburini, C., van der Zee, R., Telford, J.L., Vandenbroucke-Grauls, C.M., D’Elios, M.M., et al. (2003). Molecular mimicry between Helicobacter pylori antigens and H+, K+ --adenosine triphosphatase in human gastric autoimmunity. J Exp Med 198, 1147–1156.

Amezcua-Guerra, L.M., Rojas-Velasco, G., Brianza-Padilla, M., Vazquez-Rangel, A., Marquez-Velasco, R., Baranda-Tovar, F., Springall, R., Gonzalez-Pacheco, H., Juarez-Vicuna, Y., Tavera-Alonso, C., et al. (2020). Presence of antiphospholipid antibodies in COVID-19: case series study. Ann Rheum Dis.

Andreatta, M., and Nielsen, M. (2016). Gapped sequence alignment using artificial neural networks: application to the MHC class I system. Bioinformatics 32, 511–517.

Angileri, F., Legare, S., Marino Gammazza, A., Conway de Macario, E., Jl Macario, A., and Cappello, F. (2020a). Molecular mimicry may explain multi-organ damage in COVID-19. Autoimmun Rev 19, 102591.

Angileri, F., Legare, S., Marino Gammazza, A., Conway de Macario, E., Macario, A.J.L., and Cappello, F. (2020b). Is molecular mimicry the culprit in the autoimmune haemolytic anaemia affecting patients with COVID-19? Br J Haematol 190, e92–e93.

Bolles, M., Deming, D., Long, K., Agnihothram, S., Whitmore, A., Ferris, M., Funkhouser, W., Gralinski, L., Totura, A., Heise, M., et al. (2011). A double-inactivated severe acute respiratory syndrome coronavirus vaccine provides incomplete protection in mice and induces increased eosinophilic proinflammatory pulmonary response upon challenge. J Virol 85, 12201–12215.

Bui, H.H., Sidney, J., Dinh, K., Southwood, S., Newman, M.J., and Sette, A. (2006). Predicting population coverage of T-cell epitope-based diagnostics and vaccines. BMC Bioinformatics 7, 153.

Camacho, C., Coulouris, G., Avagyan, V., Ma, N., Papadopoulos, J., Bealer, K., and Madden, T.L. (2009). BLAST+: architecture and applications. BMC Bioinformatics 10, 421.

Cappello, F. (2020a). COVID-19 and molecular mimicry: The Columbus’ egg? J Clin Neurosci 77, 246.

Cappello, F. (2020b). Is COVID-19 a proteiform disease inducing also molecular mimicry phenomena? Cell Stress Chaperones 25, 381–382.

Cappello, F., Gammazza, A.M., Dieli, F., de, M., and Macario, A.J. (2020). Does SARS-CoV-2 Trigger Stress-InducedAutoimmunity by Molecular Mimicry? A Hypothesis. J Clin Med 9.

Chen, G., Wu, D., Guo, W., Cao, Y., Huang, D., Wang, H.W., Wang, T., Zhang, X.Y., Chen, H.L., Yu, H.J., et al. (2020). Clinical and immunological features of severe and moderate coronavirus disease 2019. Journal of Clinical Investigation 130, 2620–2629.

Chuang, Y.C., Lin, Y.S., Liu, H.S., and Yeh, T.M. (2014). Molecular mimicry between dengue virus and coagulation factors induces antibodies to inhibit thrombin activity and enhance fibrinolysis. J Virol 88, 13759–13768.

Cole, D.K., Bulek, A.M., Dolton, G., Schauenberg, A.J., Szomolay, B., Rittase, W., Trimby, A., Jothikumar, P., Fuller, A., Skowera, A., et al. (2016). Hotspot autoimmune T cell receptor binding underlies pathogen and insulin peptide cross-reactivity. J Clin Invest 126, 2191–2204.

Czub, M., Weingartl, H., Czub, S., He, R., and Cao, J. (2005). Evaluation of modified vaccinia virus Ankara based recombinant SARS vaccine in ferrets. Vaccine 23, 2273–2279.

Deming, D., Sheahan, T., Heise, M., Yount, B., Davis, N., Sims, A., Suthar, M., Harkema, J., Whitmore, A., Pickles, R., et al. (2006). Vaccine efficacy in senescent mice challenged with recombinant SARS-CoV bearing epidemic and zoonotic spike variants. PLoS Med 3, e525.

Du, L., Zhao, G., Yang, Y., Qiu, H., Wang, L., Kou, Z., Tao, X., Yu, H., Sun, S., Tseng, C.T., et al. (2014). A conformation-dependent neutralizing monoclonal antibody specifically targeting receptor-binding domain in Middle East respiratory syndrome coronavirus spike protein. J Virol 88, 7045–7053.

Eddy, S.R. (2011). Accelerated Profile HMM Searches. PLoS Comput Biol 7, e1002195.

Frick, D.N., Virdi, R.S., Vuksanovic, N., Dahal, N., and Silvaggi, N.R. (2020). Molecular Basis for ADP-Ribose Binding to the Mac1 Domain of SARS-CoV-2 nsp3. Biochemistry 59, 2608–2615.

Galeotti, C., and Bayry, J. (2020). Autoimmune and inflammatory diseases following COVID-19. Nat Rev Rheumatol 16, 413–414.

Giamarellos-Bourboulis, E.J., Netea, M.G., Rovina, N., Akinosoglou, K., Antoniadou, A., Antonakos, N., Damoraki, G., Gkavogianni, T., Adami, M.E., Katsaounou, P., et al. (2020). Complex Immune Dysregulation in COVID-19 Patients with Severe Respiratory Failure. Cell Host Microbe 27, 992–1000 e1003.

Gkoutzourelas, A., Barmakoudi, M., and Bogdanos, D.P. (2020). A Bioinformatics Analysis Reveals Novel Pathogens as Molecular Mimicry Triggers of Systemic Sclerosis. Mediterr J Rheumatol 31, 50–70.

Gourh, P., Safran, S.A., Alexander, T., Boyden, S.E., Morgan, N.D., Shah, A.A., Mayes, M.D., Doumatey, A., Bentley, A.R., Shriner, D., et al. (2020). HLA and autoantibodies define scleroderma subtypes and risk in African and European Americans and suggest a role for molecular mimicry. Proc Natl Acad Sci U S A 117, 552–562.

Greiling, T.M., Dehner, C., Chen, X., Hughes, K., Iniguez, A.J., Boccitto, M., Ruiz, D.Z., Renfroe, S.C., Vieira, S.M., Ruff, W.E., et al. (2018). Commensal orthologs of the human autoantigen Ro60 as triggers of autoimmunity in lupus. Sci Transl Med 10.

Gruber, C.N., Patel, R.S., Trachtman, R., Lepow, L., Amanat, F., Krammer, F., Wilson, K.M., Onel, K., Geanon, D., Tuballes, K., et al. (2020). Mapping Systemic Inflammation and Antibody Responses in Multisystem Inflammatory Syndrome in Children (MIS-C). Cell.

Grunewald, M.E., Chen, Y., Kuny, C., Maejima, T., Lease, R., Ferraris, D., Aikawa, M., Sullivan, C.S., Perlman, S., and Fehr, A.R. (2019). The coronavirus macrodomain is required to prevent PARP-mediated inhibition of virus replication and enhancement of IFN expression. PLoS Pathog 15, e1007756.

Hadinegoro, S.R., Arredondo-Garcia, J.L., Capeding, M.R., Deseda, C., Chotpitayasunondh, T., Dietze, R., Muhammad Ismail, H.I., Reynales, H., Limkittikul, K., Rivera-Medina, D.M., et al. (2015). Efficacy and Long-Term Safety of a Dengue Vaccine in Regions of Endemic Disease. N Engl J Med 373, 1195–1206.

Heer, C.D., Sanderson, D.J., Alhammad, Y.M.O., Schmidt, M.S., Trammell, S.A.J., Perlman, S., Cohen, M.S., Fehr, A.R., and Brenner, C. (2020). Coronavirus and PARP expression dysregulate the NAD Metabolome: a potentially actionable component of innate immunity. bioRxiv.

Heo, L., and Feig, M. (2020). Modeling of Severe Acute Respiratory Syndrome Coronavirus 2 (SARS-CoV-2) Proteins by Machine Learning and Physics-Based Refinement. bioRxiv.

Hindilerden, F., Yonal-Hindilerden, I., Akar, E., and Kart-Yasar, K. (2020). Covid-19 associated autoimmune thrombotic thrombocytopenic purpura: Report of a case. Thromb Res 195, 136–138.

Huang, X., Pearce, R., and Zhang, Y. (2020). De novo design of protein peptides to block association of the SARS-CoV-2 spike protein with human ACE2. Aging (Albany NY) 12, 11263–11276.

Hundal, J., Kiwala, S., McMichael, J., Miller, C.A., Xia, H., Wollam, A.T., Liu, C.J., Zhao, S., Feng, Y.Y., Graubert, A.P., et al. (2020). pVACtools: A Computational Toolkit to Identify and Visualize Cancer Neoantigens. Cancer Immunol Res 8, 409–420.

Iwata, H., Goettsch, C., Sharma, A., Ricchiuto, P., Bin Goh, W.W., Halu, A., Yamada, I., Yoshida, H., Hara, T., Wei, M., et al. (2016). PARP9 and PARP14 cross-regulate macrophage activation via STAT1 ADP-ribosylation. Nat Commun 7.

Jensen, C.E., Wilson, S., Thombare, A., Weiss, S., and Ma, A. (2020). Cold agglutinin syndrome as a complication of Covid-19 in two cases. Clin Infect Pract 7, 100041.

Jeoung, D., Lim, Y., Lee, E.B., Lee, S., Kim, H.Y., Lee, H., and Song, Y.W. (2004). Identification of autoantibody against poly (ADP-ribose) polymerase (PARP) fragment as a serological marker in systemic lupus erythematosus. J Autoimmun 22, 87–94.

Jespersen, M.C., Peters, B., Nielsen, M., and Marcatili, P. (2017). BepiPred-2.0: improving sequence-based B-cell epitope prediction using conformational epitopes. Nucleic Acids Res 45, W24–W29.

Ji, Q., Perchellet, A., and Goverman, J.M. (2010). Viral infection triggers central nervous system autoimmunity via activation of CD8+ T cells expressing dual TCRs. Nat Immunol 11, 628–634.

Jurtz, V., Paul, S., Andreatta, M., Marcatili, P., Peters, B., and Nielsen, M. (2017). NetMHCpan-4.0: Improved Peptide-MHC Class I Interaction Predictions Integrating Eluted Ligand and Peptide Binding Affinity Data. J Immunol 199, 3360–3368.

Kanduc, D. (2020). From Anti-SARS-CoV-2 Immune Responses to COVID-19 via Molecular Mimicry. Antibodies (Basel) 9.

Kanduc, D., and Shoenfeld, Y. (2020). Molecular mimicry between SARS-CoV-2 spike glycoprotein and mammalian proteomes: implications for the vaccine. Immunol Res 68, 310–313.

Karosiene, E., Lundegaard, C., Lund, O., and Nielsen, M. (2012). NetMHCcons: a consensus method for the major histocompatibility complex class I predictions. Immunogenetics 64, 177-186.

Katzelnick, L.C., Gresh, L., Halloran, M.E., Mercado, J.C., Kuan, G., Gordon, A., Balmaseda, A., and Harris, E. (2017). Antibody-dependent enhancement of severe dengue disease in humans. Science 358, 929–932.

Kewan, T., Almhana, F., Schwartzman, L., Daw, H., and Haddad, A. (2020). COVID-19 patient with immune thrombocytopenic purpura. Int J Lab Hematol.

Khodadoust, M.S., Olsson, N., Wagar, L.E., Haabeth, O.A., Chen, B., Swaminathan, K., Rawson, K., Liu, C.L., Steiner, D., Lund, P., et al. (2017). Antigen presentation profiling reveals recognition of lymphoma immunoglobulin neoantigens. Nature 543, 723–727.

Kim, T.S., and Shin, E.C. (2019). The activation of bystander CD8(+) T cells and their roles in viral infection. Exp Mol Med 51, 1–9.

Kim, Y., Sidney, J., Pinilla, C., Sette, A., and Peters, B. (2009). Derivation of an amino acid similarity matrix for peptide: MHC binding and its application as a Bayesian prior. BMC Bioinformatics 10, 394.

Krammer, F. (2020). SARS-CoV-2 vaccines in development. Nature.

Kudose, S., Batal, I., Santoriello, D., Xu, K., Barasch, J., Peleg, Y., Canetta, P., Ratner, L.E., Marasa, M., Gharavi, A.G., et al. (2020). Kidney Biopsy Findings in Patients with COVID-19. J Am Soc Nephrol 31, 1959–1968.

Lee, W.S., Wheatley, A.K., Kent, S.J., and DeKosky, B.J. (2020). Antibody-dependent enhancement and SARS-CoV-2 vaccines and therapies. Nat Microbiol 5, 1185–1191.

Lindsley, A.W., Schwartz, J.T., and Rothenberg, M.E. (2020). Eosinophil responses during COVID-19 infections and coronavirus vaccination. J Allergy Clin Immunol 146, 1–7.

Liu, G., Carter, B., Bricken, T., Jain, S., Viard, M., Carrington, M., and Gifford, D.K. (2020). Computationally Optimized SARS-CoV-2 MHC Class I and II Vaccine Formulations Predicted to Target Human Haplotype Distributions. Cell Syst 11, 131–144 e136.

Liu, L., Wei, Q., Lin, Q., Fang, J., Wang, H., Kwok, H., Tang, H., Nishiura, K., Peng, J., Tan, Z., et al. (2019). Anti-spike IgG causes severe acute lung injury by skewing macrophage responses during acute SARS-CoV infection. JCI Insight 4.

Lopez, C., Kim, J., Pandey, A., Huang, T., and DeLoughery, T.G. (2020). Simultaneous onset of COVID-19 and autoimmune haemolytic anaemia. Br J Haematol 190, 31–32.

Lucchese, G., and Floel, A. (2020a). Molecular mimicry between SARS-CoV-2 and respiratory pacemaker neurons. Autoimmun Rev 19, 102556.

Lucchese, G., and Floel, A. (2020b). SARS-CoV-2 and Guillain-Barre syndrome: molecular mimicry with human heat shock proteins as potential pathogenic mechanism. Cell Stress Chaperones 25, 731–735.

Luo, F., Liao, F.L., Wang, H., Tang, H.B., Yang, Z.Q., and Hou, W. (2018a). Evaluation of Antibody-Dependent Enhancement of SARS-CoV Infection in Rhesus Macaques Immunized with an Inactivated SARS-CoV Vaccine. Virol Sin 33, 201–204.

Luo, G., Ambati, A., Lin, L., Bonvalet, M., Partinen, M., Ji, X., Maecker, H.T., and Mignot, E.J. (2018b). Autoimmunity to hypocretin and molecular mimicry to flu in type 1 narcolepsy. Proc Natl Acad Sci U S A 115, E12323–E12332.

Magnan, C.N., and Baldi, P. (2014). SSpro/ACCpro 5: almost perfect prediction of protein secondary structure and relative solvent accessibility using profiles, machine learning and structural similarity. Bioinformatics 30, 2592–2597.

Mallapaty, S., and Ledford, H. (2020). COVID-vaccine results are on the way - and scientists’ concerns are growing. Nature 586, 16–17.

Mandal, K., Jahangiri, M., and Xu, Q.B. (2004). Autoimmunity to heat shock proteins in atherosclerosis. Autoimmunity Reviews 3, 121–127.

Marino Gammazza, A., Legare, S., Lo Bosco, G., Fucarino, A., Angileri, F., Conway de Macario, E., Macario, A.J., and Cappello, F. (2020). Human molecular chaperones share with SARS-CoV-2 antigenic epitopes potentially capable of eliciting autoimmunity against endothelial cells: possible role of molecular mimicry in COVID-19. Cell Stress Chaperones 25, 737–741.

Mathew, D., Giles, J.R., Baxter, A.E., Oldridge, D.A., Greenplate, A.R., Wu, J.E., Alanio, C., Kuri-Cervantes, L., Pampena, M.B., D’Andrea, K., et al. (2020). Deep immune profiling of COVID-19 patients reveals distinct immunotypes with therapeutic implications. Science 369.

Matzaraki, V., Kumar, V., Wijmenga, C., and Zhernakova, A. (2017). The MHC locus and genetic susceptibility to autoimmune and infectious diseases. Genome Biol 18, 76.

McCoy, L., Tsunoda, I., and Fujinami, R.S. (2006). Multiple sclerosis and virus induced immune responses: autoimmunity can be primed by molecular mimicry and augmented by bystander activation. Autoimmunity 39, 9–19.

Megremis, S., Walker, T.D.J., He, X., Ollier, W.E.R., Chinoy, H., Hampson, L., Hampson, I., and Lamb, J.A. (2020). Antibodies against immunogenic epitopes with high sequence identity to SARS-CoV-2 in patients with autoimmune dermatomyositis. Ann Rheum Dis 79, 1383–1386.

Mehan, W.A., Yoon, B.C., Lang, M., Li, M.D., Rincon, S., and Buch, K. (2020). Paraspinal Myositis in Patients with COVID-19 Infection. AJNR Am J Neuroradiol 41, 1949–1952.

Merrill, J.T., Erkan, D., Winakur, J., and James, J.A. (2020). Emerging evidence of a COVID-19 thrombotic syndrome has treatment implications. Nat Rev Rheumatol 16, 581–589.

Miller, S.D., Vanderlugt, C.L., Begolka, W.S., Pao, W., Yauch, R.L., Neville, K.L., Katz-Levy, Y., Carrizosa, A., and Kim, B.S. (1997). Persistent infection with Theiler’s virus leads to CNS autoimmunity via epitope spreading. Nat Med 3, 1133–1136.

Neuman, B.W., Joseph, J.S., Saikatendu, K.S., Serrano, P., Chatterjee, A., Johnson, M.A., Liao, L., Klaus, J.P., Yates, J.R., 3rd, Wuthrich, K., et al. (2008). Proteomics analysis unravels the functional repertoire of coronavirus nonstructural protein 3. J Virol 82, 5279–5294.

Nielsen, M., and Andreatta, M. (2017). NNAlign: a platform to construct and evaluate artificial neural network models of receptor-ligand interactions. Nucleic Acids Res 45, W344–W349.

Nielsen, M., Lundegaard, C., and Lund, O. (2007). Prediction of MHC class II binding affinity using SMM-align, a novel stabilization matrix alignment method. BMC Bioinformatics 8, 238.

O’Donnell, T.J., Rubinsteyn, A., and Laserson, U. (2020). MHCflurry 2.0: Improved Pan-Allele Prediction of MHC Class I-Presented Peptides by Incorporating Antigen Processing. Cell Syst 11, 42–48 e47.

Ong, E., Wong, M.U., Huffman, A., and He, Y. (2020). COVID-19 Coronavirus Vaccine Design Using Reverse Vaccinology and Machine Learning. Front Immunol 11, 1581.

Otsuka, R., and Seino, K.I. (2020). Macrophage activation syndrome and COVID-19. Inflamm Regen 40, 19.

Ottaviani, D., Boso, F., Tranquillini, E., Gapeni, I., Pedrotti, G., Cozzio, S., Guarrera, G.M., and Giometto, B. (2020). Early Guillain-Barre syndrome in coronavirus disease 2019 (COVID-19): a case report from an Italian COVID-hospital. Neurol Sci 41, 1351–1354.

Petersen, J., Ciacchi, L., Tran, M.T., Loh, K.L., Kooy-Winkelaar, Y., Croft, N.P., Hardy, M.Y., Chen, Z., McCluskey, J., Anderson, R.P., et al. (2020). T cell receptor cross-reactivity between gliadin and bacterial peptides in celiac disease. Nat Struct Mol Biol 27, 49–61.

Phillips, N., Cyranoski, D., and Mallapaty, S. (2020). A leading coronavirus vaccine trial is on hold: scientists react. Nature.

Polymeros, D., Tsiamoulos, Z.P., Koutsoumpas, A.L., Smyk, D.S., Mytilinaiou, M.G., Triantafyllou, K., Bogdanos, D.P., and Ladas, S.D. (2014). Bioinformatic and immunological analysis reveals lack of support for measles virus related mimicry in Crohn’s disease. BMC Med 12,139.

Qin, C., Zhou, L., Hu, Z., Zhang, S., Yang, S., Tao, Y., Xie, C., Ma, K., Shang, K., Wang, W., et al. (2020). Dysregulation of Immune Response in Patients With Coronavirus 2019 (COVID-19) in Wuhan, China. Clin Infect Dis 71, 762–768.

Reynisson, B., Barra, C., Kaabinejadian, S., Hildebrand, W.H., Peters, B., and Nielsen, M. (2020). Improved Prediction of MHC II Antigen Presentation through Integration and Motif Deconvolution of Mass Spectrometry MHC Eluted Ligand Data. J Proteome Res 19, 2304–2315.

Riley, T.P., Hellman, L.M., Gee, M.H., Mendoza, J.L., Alonso, J.A., Foley, K.C., Nishimura, M.I., Vander Kooi, C.W., Garcia, K.C., and Baker, B.M. (2018). T cell receptor cross-reactivity expanded by dramatic peptide-MHC adaptability. Nat Chem Biol 14, 934–942.

Rodriguez, Y., Novelli, L., Rojas, M., De Santis, M., Acosta-Ampudia, Y., Monsalve, D.M., Ramirez-Santana, C., Costanzo, A., Ridgway, W.M., Ansari, A.A., et al. (2020). Autoinflammatory and autoimmune conditions at the crossroad of COVID-19. J Autoimmun, 102506.

Rodriguez, Y., Rojas, M., Pacheco, Y., Acosta-Ampudia, Y., Ramirez-Santana, C., Monsalve, D.M., Gershwin, M.E., and Anaya, J.M. (2018). Guillain-Barre syndrome, transverse myelitis and infectious diseases. Cell Mol Immunol 15, 547–562.

Rojas, M., Restrepo-Jimenez, P., Monsalve, D.M., Pacheco, Y., Acosta-Ampudia, Y., Ramirez-Santana, C., Leung, P.S.C., Ansari, A.A., Gershwin, M.E., and Anaya, J.M. (2018). Molecular mimicry and autoimmunity. J Autoimmun 95, 100–123.

Ruff, W.E., Greiling, T.M., and Kriegel, M.A. (2020). Host-microbiota interactions in immune-mediated diseases. Nat Rev Microbiol 18, 521–538.

Schiaffino, M.T., Di Natale, M., Garcia-Martinez, E., Navarro, J., Munoz-Blanco, J.L., Demelo-Rodriguez, P., and Sanchez-Mateos, P. (2020). Immunoserologic Detection and Diagnostic Relevance of Cross-Reactive Autoantibodies in Coronavirus Disease 2019 Patients. J Infect Dis 222, 1439–1443.

Sedaghat, Z., and Karimi, N. (2020). Guillain Barre syndrome associated with COVID-19 infection: A case report. J Clin Neurosci 76, 233–235.

Seitz-Polski, B., Debiec, H., Rousseau, A., Dahan, K., Zaghrini, C., Payre, C., Esnault, V.L.M., Lambeau, G., and Ronco, P. (2018). Phospholipase A2 Receptor 1 Epitope Spreading at Baseline Predicts Reduced Likelihood of Remission of Membranous Nephropathy. J Am Soc Nephrol 29, 401–408.

Shah, S., Danda, D., Kavadichanda, C., Das, S., Adarsh, M.B., and Negi, V.S. (2020). Autoimmune and rheumatic musculoskeletal diseases as a consequence of SARS-CoV-2 infection and its treatment. Rheumatol Int 40, 1539–1554.

Shao, X.M., Bhattacharya, R., Huang, J., Sivakumar, I.K.A., Tokheim, C., Zheng, L., Hirsch, D., Kaminow, B., Omdahl, A., Bonsack, M., et al. (2020). High-Throughput Prediction of MHC Class I and II Neoantigens with MHCnuggets. Cancer Immunol Res 8, 396–408.

Sokolove, J., Bromberg, R., Deane, K.D., Lahey, L.J., Derber, L.A., Chandra, P.E., Edison, J.D., Gilliland, W.R., Tibshirani, R.J., Norris, J.M., et al. (2012). Autoantibody epitope spreading in the pre-clinical phase predicts progression to rheumatoid arthritis. PLoS One 7, e35296.

Steinegger, M., and Soding, J. (2017). MMseqs2 enables sensitive protein sequence searching for the analysis of massive data sets. Nat Biotechnol 35, 1026–1028.

Su, H., Yang, M., Wan, C., Yi, L.X., Tang, F., Zhu, H.Y., Yi, F., Yang, H.C., Fogo, A.B., Nie, X., et al. (2020). Renal histopathological analysis of 26 postmortem findings of patients with COVID-19 in China. Kidney Int 98, 219–227.

Suso, A.S., Mon, C., Alonso, I.O., Romo, K.G., Juarez, R.C., Ramirez, C.L., Sanchez, M.S., Valdivia, V.M., Librero, M.O., Pala, A.O., et al. (2020). IgA Vasculitis with Nephritis (Henoch-Schonlein purpura) in a COVID-19 patient. Kidney Int Rep.

Suzek, B.E., Wang, Y., Huang, H., McGarvey, P.B., Wu, C.H., and UniProt, C. (2015). UniRef clusters: a comprehensive and scalable alternative for improving sequence similarity searches. Bioinformatics 31, 926–932.

Talotta, R., and Robertson, E. (2020). Autoimmunity as the comet tail of COVID-19 pandemic. World J Clin Cases 8, 3621–3644.

Tandon, R., Sharma, M., Chandrashekhar, Y., Kotb, M., Yacoub, M.H., and Narula, J. (2013). Revisiting the pathogenesis of rheumatic fever and carditis. Nat Rev Cardiol 10, 171–177.

Tien, M.Z., Meyer, A.G., Sydykova, D.K., Spielman, S.J., and Wilke, C.O. (2013). Maximum allowed solvent accessibilites of residues in proteins. PLoS One 8, e80635.

Tsao, H.S., Chason, H.M., and Fearon, D.M. (2020). Immune Thrombocytopenia (ITP) in a Pediatric Patient Positive for SARS-CoV-2. Pediatrics 146.

Tseng, C.T., Sbrana, E., Iwata-Yoshikawa, N., Newman, P.C., Garron, T., Atmar, R.L., Peters, C.J., and Couch, R.B. (2012). Immunization with SARS coronavirus vaccines leads to pulmonary immunopathology on challenge with the SARS virus. PLoS One 7, e35421.

Uppal, N.N., Kello, N., Shah, H.H., Khanin, Y., De Oleo, I.R., Epstein, E., Sharma, P., Larsen, C.P., Bijol, V., and Jhaveri, K.D. (2020). De Novo ANCA-associated Vasculitis with Glomerulonephritis in COVID-19. Kidney Int Rep.

Venigalla, S.S.K., Premakumar, S., and Janakiraman, V. (2020). A possible role for autoimmunity through molecular mimicry in alphavirus mediated arthritis. Sci Rep 10, 938.

Verdoni, L., Mazza, A., Gervasoni, A., Martelli, L., Ruggeri, M., Ciuffreda, M., Bonanomi, E., and D’Antiga, L. (2020). An outbreak of severe Kawasaki-like disease at the Italian epicentre of the SARS-CoV-2 epidemic: an observational cohort study. Lancet 395, 1771–1778.

Vita, R., Mahajan, S., Overton, J.A., Dhanda, S.K., Martini, S., Cantrell, J.R., Wheeler, D.K., Sette, A., and Peters, B. (2019). The Immune Epitope Database (IEDB): 2018 update. Nucleic Acids Res 47, D339–D343.

Wan, Y., Shang, J., Sun, S., Tai, W., Chen, J., Geng, Q., He, L., Chen, Y., Wu, J., Shi, Z., et al. (2020). Molecular Mechanism for Antibody-Dependent Enhancement of Coronavirus Entry. J Virol 94.

Waterhouse, A., Bertoni, M., Bienert, S., Studer, G., Tauriello, G., Gumienny, R., Heer, F.T., de Beer, T.A.P., Rempfer, C., Bordoli, L., et al. (2018). SWISS-MODEL: homology modelling of protein structures and complexes. Nucleic Acids Res 46, W296–W303.

Webb, B., and Sali, A. (2016). Comparative Protein Structure Modeling Using MODELLER. Curr Protoc Bioinformatics 54, 5 6 1–5 6 37.

Whalley, T., Dolton, G., Brown, P.E., Wall, A., Wooldridge, L., van den Berg, H., Fuller, A., Hopkins, J.R., Crowther, M.D., Attaf, M., et al. (2020). GPU-Accelerated Discovery of Pathogen-Derived Molecular Mimics of a T-Cell Insulin Epitope. Front Immunol 11, 296.

Whittaker, E., Bamford, A., Kenny, J., Kaforou, M., Jones, C.E., Shah, P., Ramnarayan, P., Fraisse, A., Miller, O., Davies, P., et al. (2020). Clinical Characteristics of 58 Children With a Pediatric Inflammatory Multisystem Syndrome Temporally Associated With SARS-CoV-2. JAMA 324, 259–269.

Wieczorek, M., Abualrous, E.T., Sticht, J., Alvaro-Benito, M., Stolzenberg, S., Noe, F., and Freund, C. (2017). Major Histocompatibility Complex (MHC) Class I and MHC Class II Proteins: Conformational Plasticity in Antigen Presentation. Front Immunol 8, 292.

Wucherpfennig, K.W., and Strominger, J.L. (1995). Molecular Mimicry in T-Cell-Mediated Autoimmunity - Viral Peptides Activate Human T-Cell Clones Specific for Myelin Basic-Protein. Cell 80, 695–705.

Yasui, F., Kai, C., Kitabatake, M., Inoue, S., Yoneda, M., Yokochi, S., Kase, R., Sekiguchi, S., Morita, K., Hishima, T., et al. (2008). Prior immunization with severe acute respiratory syndrome (SARS)-associated coronavirus (SARS-CoV) nucleocapsid protein causes severe pneumonia in mice infected with SARS-CoV. J Immunol 181, 6337–6348.

Yin, J., Sternes, P.R., Wang, M., Song, J., Morrison, M., Li, T., Zhou, L., Wu, X., He, F., Zhu, J., et al. (2020). Shotgun metagenomics reveals an enrichment of potentially cross-reactive bacterial epitopes in ankylosing spondylitis patients, as well as the effects of TNFi therapy upon microbiome composition. Ann Rheum Dis 79, 132–140.

Yip, M.S., Leung, N.H., Cheung, C.Y., Li, P.H., Lee, H.H., Daeron, M., Peiris, J.S., Bruzzone, R., and Jaume, M. (2014). Antibody-dependent infection of human macrophages by severe acute respiratory syndrome coronavirus. Virol J 11, 82.

Zhang, H., Charmchi, Z., Seidman, R.J., Anziska, Y., Velayudhan, V., and Perk, J. (2020a). COVID-19-associated myositis with severe proximal and bulbar weakness. Muscle Nerve 62, E57–E60.

Zhang, H., Lund, O., and Nielsen, M. (2009). The PickPocket method for predicting binding specificities for receptors based on receptor pocket similarities: application to MHC-peptide binding. Bioinformatics 25, 1293–1299.

Zhang, Y., Xiao, M., Zhang, S., Xia, P., Cao, W., Jiang, W., Chen, H., Ding, X., Zhao, H., Zhang, H., et al. (2020b). Coagulopathy and Antiphospholipid Antibodies in Patients with Covid-19. N Engl J Med 382, e38.

Zhao, Q., Meng, M., Kumar, R., Wu, Y., Huang, J., Deng, Y., Weng, Z., and Yang, L. (2020). Lymphopenia is associated with severe coronavirus disease 2019 (COVID-19) infections: A systemic review and meta-analysis. Int J Infect Dis 96, 131–135.

Zhao, Z.S., Granucci, F., Yeh, L., Schaffer, P.A., and Cantor, H. (1998). Molecular mimicry by herpes simplex virus-type 1: autoimmune disease after viral infection. Science 279, 1344–1347.

Zhu, J., Quyyumi, A.A., Rott, D., Csako, G., Wu, H., Halcox, J., and Epstein, S.E. (2001). Antibodies to human heat-shock protein 60 are associated with the presence and severity of coronary artery disease: evidence for an autoimmune component of atherogenesis. Circulation 103, 1071–1075.

Zulfiqar, A.A., Lorenzo-Villalba, N., Hassler, P., and Andres, E. (2020). Immune Thrombocytopenic Purpura in a Patient with Covid-19. N Engl J Med 382, e43.

